# Pan-Cancer landscape of protein activities identifies drivers of signalling dysregulation and patient survival

**DOI:** 10.1101/2021.06.09.447741

**Authors:** Abel Sousa, Aurelien Dugourd, Danish Memon, Borgthor Petursson, Evangelia Petsalaki, Julio Saez-Rodriguez, Pedro Beltrao

## Abstract

Genetic alterations in cancer cells trigger oncogenic transformation, a process largely mediated by the dysregulation of kinase and transcription factor (TF) activities. While the mutational profiles of thousands of tumours has been extensively characterized, the measurements of protein activities has been technically limited until recently. We compiled public data of matched genomics and (phospho)proteomics measurements for 1,110 tumours and 77 cell lines that we used to estimate activity changes in 218 kinases and 292 TFs. Kinase activities are, on average, not strongly determined by protein abundance but rather by their phosphorylation state while the reverse is more common for TFs. Co-regulation of kinase and TF activities reflects previously known regulatory relationships and allows us to dissect genetic drivers of signalling changes in cancer. Loss-of-function mutation is not often associated with dysregulation of downstream targets, suggesting frequent compensatory mechanisms. Finally, we identified the activities most differentially regulated in cancer subtypes and showed how these can be linked to differences in patient survival. Our results provide broad insights into dysregulation of protein activities in cancer and their contribution to disease severity.

## Introduction

Cancer is a highly heterogeneous disease that is generally caused by the acquisition of somatic genomic alterations, including single nucleotide variants (SNVs), gene copy-number variations (CNVs) and large chromosomal rearrangements (Pleasance et al. 2010; Beroukhim et al. 2010; ICGC/TCGA Pan-Cancer Analysis of Whole Genomes Consortium 2020). The Cancer Genome Atlas (TCGA) has led to an in-depth characterization of the genomic alterations of more than 10,000 tumours from 33 cancer types (Hoadley et al. 2018; Ding et al. 2018). However, mutations in key driver genes are just the first steps of a cascade of events that culminate in tumour formation and cancer. These mutations generate the genetic diversity that promotes the acquisition of multiple cancer hallmarks, including chronic proliferation, resistance to cell death and tissue invasion and metastasis (Hanahan and Weinberg 2011). An understanding of the molecular mechanisms that underpin the development of cancer is critical in order to study cancer biology and to develop therapies.

While somatic alterations and gene expression changes across tumours have been extensively studied, key driver genomic changes in cancer are thought to result in changes in cell signalling including the misregulation of protein kinases and transcription factors (Yaffe 2019; Blume-Jensen and Hunter 2001). As an example, about 40% of melanomas contain the V600E activating mutation in the BRAF kinase, resulting in constitutive signalling through the Raf to mitogen-activated protein kinase (MAPK) pathway and increased cellular proliferation (Davies and Samuels 2010). Likewise, aberrant transcription factors (TFs) activities is a key feature of cancer cells (Garcia-Alonso et al. 2018). TFs are commonly dysregulated due to genomic alterations in their sequences or in upstream signalling regulatory proteins (Oliner et al. 1992; Ohh et al. 2000). Because of their role as signalling effectors, aberrant kinase signalling may dysregulate the activities of TFs and alter the expression of their target genes. Consequently, kinases and TFs often accumulate cancer driver mutations, such as TP53 (Rivlin et al. 2011) and KRAS (Wang et al. 2015), and are the targets of anti-cancer drugs (Bhagwat and Vakoc 2015; Bhullar et al. 2018).

Due to technical limitations, the study of protein signalling activities has been for many years limited primarily to the study of a few key signalling proteins at a time using antibodies, which was recently expanded to a few hundreds via the use of reverse-phase protein arrays (RPPA) (J. Li et al. 2013). The Clinical Proteomic Tumour Analysis Consortium (CPTAC) has revolutionized the study of cancer proteomes, including proteins and respective post-translational modifications (PTMs), through the application of Mass Spectrometry (MS)-based proteomics (Bing Zhang et al. 2019). MS-based proteomic profiling of human cancers has the potential to uncover molecular insights that might be otherwise missed by genomics- and transcriptomics-driven cancer research. CPTAC enabled to (i) identify additional cancer molecular subtypes (Mun et al. 2019; Gao et al. 2019), (ii) find that changes at the genomic and transcriptomic level are often buffered at the proteomic level (Mertins et al. 2016; Bing Zhang et al. 2014; Gonçalves et al. 2017; Sousa et al. 2019) and (iii) uncover dysregulated signalling pathways by phosphoproteomics data integration (Clark et al. 2019).

In efforts to find novel therapeutic opportunities from kinase and TF oncogenic signalling, it is crucial to understand how the activities of these key signalling proteins are changing across tumours. Previous studies found that TF mutations were correlated with transcriptional dysregulation in cancer cell lines and primary tumours, and that TF activities can act as predictors of sensitivity to anti-cancer drugs (Garcia-Alonso et al. 2018). Similar results were found regarding the impact of oncogenic mutations on kinase signalling. However, these studies were focused on few kinases and cancer types (Guo et al. 2008; Guha et al. 2008; Creixell et al. 2015; Lundby et al. 2019). Despite all of these efforts, a systematic Pan-Cancer analysis of the regulation of kinase and TF activities across tumours is still lacking.

In this study, we mined multi-omics datasets from patient tumours and cancer cell lines to study the regulation of kinases and TFs across tumour types. We estimated the activities of TFs and kinases from the gene expression levels and phosphorylation changes of their targets, deriving activity profiles of 292 TFs and 218 kinases across 1,110 primary tumors from TCGA and CPTAC and 77 cancer cell lines. We used these kinase and TF activities to study the principles of regulation of these signalling proteins by mutations, changes in abundance or phosphorylation. We show how their patterns of activity co-regulation reflect underlying signalling relationships and we identify the signalling molecules that show high degree of regulation in each tumour type. Finally, we show how these TF/kinase activities can be predictive of differential survival across patients. The profile of protein activities across over 1000 patient samples serves as a resource to study the misregulation of signalling across different tumour types.

## Results

### Standardized multi-omics pan-cancer dataset

To study the regulation of protein activities of cancer cells, we compiled and standardized multi-omics datasets made available by the CPTAC consortium (**Figure 1A; Methods**). These datasets comprised of cancer patient samples with matched somatic mutations, gene copy number variation (CNV), mRNA expression, protein abundance, phosphorylation and clinical data from 9 tissues: breast (Mertins et al. 2016; Cancer Genome Atlas Network 2012b), brain (Petralia et al. 2020), colorectal (Bing Zhang et al. 2014; Cancer Genome Atlas Network 2012a; Vasaikar et al. 2019), ovarian (H. Zhang et al. 2016; Cancer Genome Atlas Research Network 2011), liver (Gao et al. 2019), kidney (Clark et al. 2019), uterus (Dou et al. 2020), lung (Gillette et al. 2020) and stomach (Mun et al. 2019). In addition, we collected data for breast (Lapek et al. 2017; Lawrence et al. 2015) and colorectal (Roumeliotis et al. 2017) cancer cell lines, for which multi-omics data were available (**Figure 1A; Methods**). In summary, the assembled data provides the opportunity to build an integrated picture of the cancer genome, transcriptome and (phospho)proteome, with 1008 samples (932 tumours and 76 cell lines) matching all data types available per dataset.

**Figure 1.**
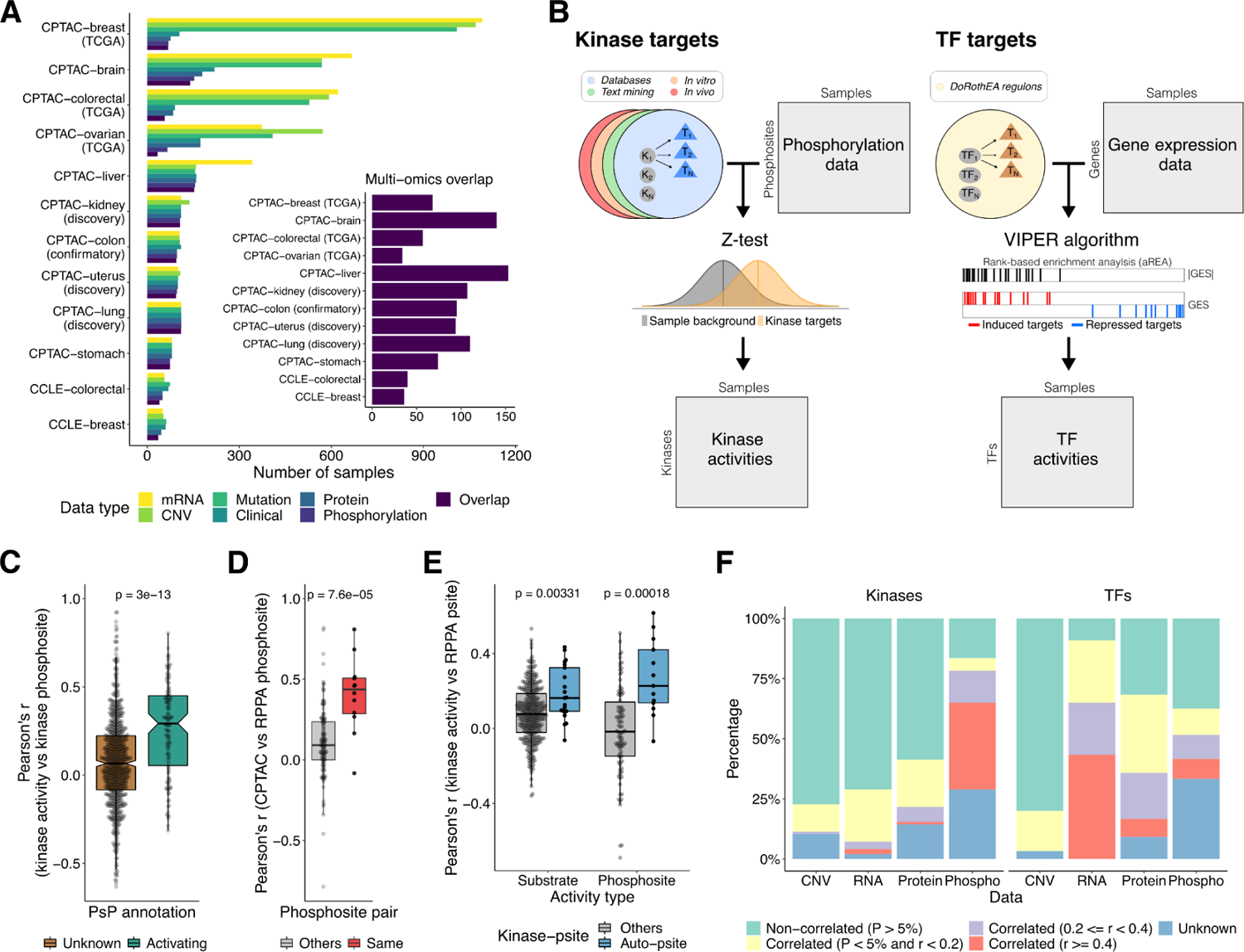
Multi-omics atlas and inference of protein activities. **(A)** Number of samples by cancer dataset and data type. **(B)** Schematic representation of kinase and TF activity inference. GES, gene expression signature. **(C)** Comparison of the Pearson’s correlation distributions between the kinase activities and the quantifications of phosphosites (log2 fold-changes) that mapped to the same kinase, with (n = 126) and without (n = 793) annotation (activating) in PhosphoSitePlus. A P-value from a Wilcoxon rank sum test is shown. **(D)** Pearson’s correlation between the CPTAC MS-based and the TCGA RPPA-based phosphosite quantifications, for the same phosphosite pair (n = 12) and others (n = 132). A P-value from a Wilcoxon rank sum test is shown. **(E)** Comparison of the Pearson’s correlation between the RPPA phosphosites and the kinase activities, for kinase-phosphosite pairs mapping to the same kinase (auto-phosphosite) and other pairs. The activities were calculated using the kinase substrates (n = 336 and n = 21) and the kinase regulatory phosphosites (n = 94 and n = 13) (**Methods**). The P-values from Wilcoxon rank sum tests are shown. **(F)** Percentage of kinases and TFs significantly and not significantly correlated with the corresponding CNV, RNA, protein and phosphorylation levels. The proteins without correlations due to lack of data or reduced number of samples (n < 10) were labeled as unknown (blue).

We first calculated correlations between each protein and phosphosites that mapped to the same protein, across up to 1008 samples. Across all pairs there is a correlation of 0.49 (p-value < 2.2×10e-16), with an average protein-phosphosite correlation of 0.39 (**Figures S1A-S1B**), in agreement with previous studies (Arshad et al. 2019). This result shows that phosphorylation levels are, to some extent, confounded by the corresponding protein abundance (Wu et al. 2011). To be able to focus on phosphorylation changes that are not driven primarily by protein abundance differences, we regressed-out matched protein abundance from the phosphorylation data in our compiled dataset (**Figures S1A-S1B; Methods**).

### Landscape of protein activities in cancer

The genomics characterization of tumour samples has so far been primarily focused on stratifying samples by their mutational profiles or changes in abundance of specific bio-molecules such as transcripts, protein or phosphorylation states. We and others have shown that changes in phosphorylation and gene expression levels can be used to infer the activation states of protein kinases and TFs (Ochoa et al. 2016; Garcia-Alonso et al. 2018; Casado et al. 2013; Hernandez-Armenta et al. 2017). Based on these methods we set out to define the landscape of kinase/TF activity patterns across these tumour samples.

The kinase activities were estimated from the protein abundance-corrected phosphorylation data using a z-test (Hernandez-Armenta et al. 2017) (**Figure 1B; Methods**). Briefly, the activity of a given kinase in a sample is estimated by comparing the changes in phosphorylation of its substrates with changes of all other phosphosites. Similarly, the activation state of TFs were inferred from the changes in gene expression of their known transcriptional targets using the DoRothEA regulons (Garcia-Alonso et al. 2019) coupled with the VIPER algorithm (Alvarez et al. 2016) (**Figure 1B; Methods**). In total, we estimated the activities of 292 TFs across 1,187 cancer samples (1,110 primary tumours and 77 cell lines) (**Table S1**). For the estimation of kinase activities, we evaluated different lists of kinase substrates from repositories (e.g. PhosphositePlus (Hornbeck et al. 2015)), computational text mining (Bachman, Gyori, and Sorger 2019), kinase inhibitor experiments (Hijazi et al. 2020) or phosphorylation of cell extracts (Sugiyama, Imamura, and Ishihama 2019) (**Figures S2A-S2B**). We tested each list in a compilation of phosphoproteomic experiments where kinase regulation is known (Hernandez-Armenta et al. 2017) (**Methods**) keeping those from repositories and text-mining as the most accurate (**Figures S3A-S3B**). After applying this approach we inferred the activities of 218 kinases across 980 samples (930 tumours and 50 cell lines) (**Table S1, Methods**).

For some kinases, there are phosphosites within the kinase itself that are known to activate or inhibit it. As a validation, we correlated the estimated activity scores with the quantifications of activating phosphosites, finding the expected higher correlation when compared with phosphosites without annotation (**Figure 1C**). A similar trend was observed when excluding the kinase auto-regulatory phosphosites before re-estimating the activities (**Figure S3C).** Finally, we benchmarked the kinase activity scores using reverse phase protein array (RPPA) data from the TCGA program. We first evaluated the agreement between the MS-based and the RPPA-based phosphosite quantifications, and found that phosphosite pairs corresponding to the same phosphosite show higher correlations than random pairs (**Figure 1D**). Then, we found that the RPPA phosphosites correlate significantly better with the activity of kinase bearing the phosphosites than with other kinase activities (**Figure 1E**).

The activity profiles of kinase and TFs across a large number of samples allows us to ask how these activities are themselves regulated. We first selected 99 kinases and 120 TFs that are strongly regulated in at least 5% of all samples (**Figure S3D**). We then correlated these activities with changes in gene copy number (CNV), mRNA and protein levels or changes in phosphorylation levels of the respective protein (**Table S2**). We observed that 55% of kinase activities correlated with their phosphorylation state and only 27% correlated with changes in protein abundance (**Figure 1F**). Contrary to this, TF activities are most often correlated with changes in abundance of the TF, as measured by RNA (91%) or protein (59%), with fewer cases of significant correlations with phosphorylation levels (29%) (**Figure 1F**). TF phosphosites predicted to be important for function (Ochoa et al. 2020) are more likely to show significant correlations with the TF activity (**Figure S3E**).

Overall, these results showed that our kinase activity estimates are likely to capture kinase regulatory events across different tumour types, and therefore the usefulness of our multi-omics atlas to study kinase signalling in cancer.

### Impact of genetic variation on protein abundance and activities

The large number of cancer samples in this study constitutes a resource to measure the effects of genetic alterations, i.e. somatic mutations and CNVs, on protein abundances and activities. We first set out to assess the effects of CNVs on the mRNA and protein abundances. Similarly to our previous reports, the CNVs showed a stronger correlation with the mRNA than with the protein levels (**Figures S4A-S4B**), highlighting mechanisms of post-transcriptional control and gene dosage buffering at the protein level (Sousa et al. 2019; Gonçalves et al. 2017). We then extended the analysis to globally assess the effects of mutations (**Methods**), and we found that proteins carrying loss-of-function (LoF) alterations, including frameshift, nonsense, splice site and stop codon loss, caused on average a significant decrease in protein abundance. This was not observed with in-frame and missense mutations (**Figure S4C**). To validate the decrease of protein abundance for LoF mutations, we confirmed that this was also recapitulated in a proteomic dataset with 125 cancer cell lines (CCLs) from the NCI60 and CRC65 panels (Frejno et al. 2020) (**Figure S4D; Methods**). These observations confirm that the genetic alterations are often recapilated at the protein level as captured by the MS data.

We next looked at the impact of genetic alterations on TF and kinase activity estimates. On average we did not observe reduced activity for proteins carrying different types of mutations in the tumour samples (**Figure S5A**), with only a very modest average decrease in activities for frameshift mutations found in the cell line data (**Figure S5B**). This observation did not depend on the degree of predicted deleterious impact of the mutations (**Figure S5C-S5D)** nor on the purity of the tumour samples (**Figure S5E)**. To further characterize this unexpected result, we focused on highly mutated cancer genes. As an example, we investigated the impact of the BRAF^V600E^ mutation on the predicted activities of proteins from the MAPK/ERK signaling transduction pathway (**Methods**). Surprisingly, across all samples BRAF^V600E^ mutations were not significantly associated with changes in activity of key pathway components, including BRAF itself, MAPK1, MAPK3, MAP2K1 and MAP2K2 (**Figure S5F**). Instead, we found that CDK1 and CDK7 were more active in samples carrying the mutation (FDR < 5%) (**Figure S5G**). This suggests that samples carrying a BRAF^V600E^ mutation will often have kinase activity levels that have adapted to the mutational state, likely having increased proliferation, as indicated by the CDK1 levels but not a higher level of activation of the pathway.

We then extended the analysis by systematically associating the activity of kinases and TFs with the recurrent mutational status of any given gene mutated in at least 5 tumour samples (**Methods**). As seen for the BRAF example, we didn’t observe any case where recurrent mutation of the kinase itself was associated with a significant change in its activity as measured by the phosphorylation of its substrates. This indicates that there is significant adaptation of the signalling state of the cell after mutations. On the other hand, we found 193 significant associations (FDR < 5%) between mutations in other genes and changes in kinase activity levels (**Figure 2A; Table S3**). For example, samples with mutations on STK11 (serine/threonine kinase 11) don’t show a pronounced change in activity of STK11 substrates but have decreased activity for PRKACA kinase, a known activator of STK11 (FDR = 9.6e-5; combined string network weight = 0.94) (**Figure 2C**). Other examples include increased activity for CDK1 and MAPK13 in samples with mutations in TP53, and for AKT3 when PTEN is mutated (FDR < 5%) (**Figure 2C**). Unlike for kinases, we found several cases where the mutation of a TF was associated with a change in its own activity as is the case for mutations in TP53, GATA6, SREBF2 and EBF1 (FDR < 5%) (**Figures 2B - inner plot, S6; Table S3**). In addition we found 11,128 significant associations between a mutated gene and a changed TF activity (FDR < 5%) (1,087 for FDR < 1%) (**Figure 2B - outer plot; Table S3**), including increased activity for E2F4 and TFDP1 coupled with TP53 mutation (**Figure 2D**).

**Figure 2.**
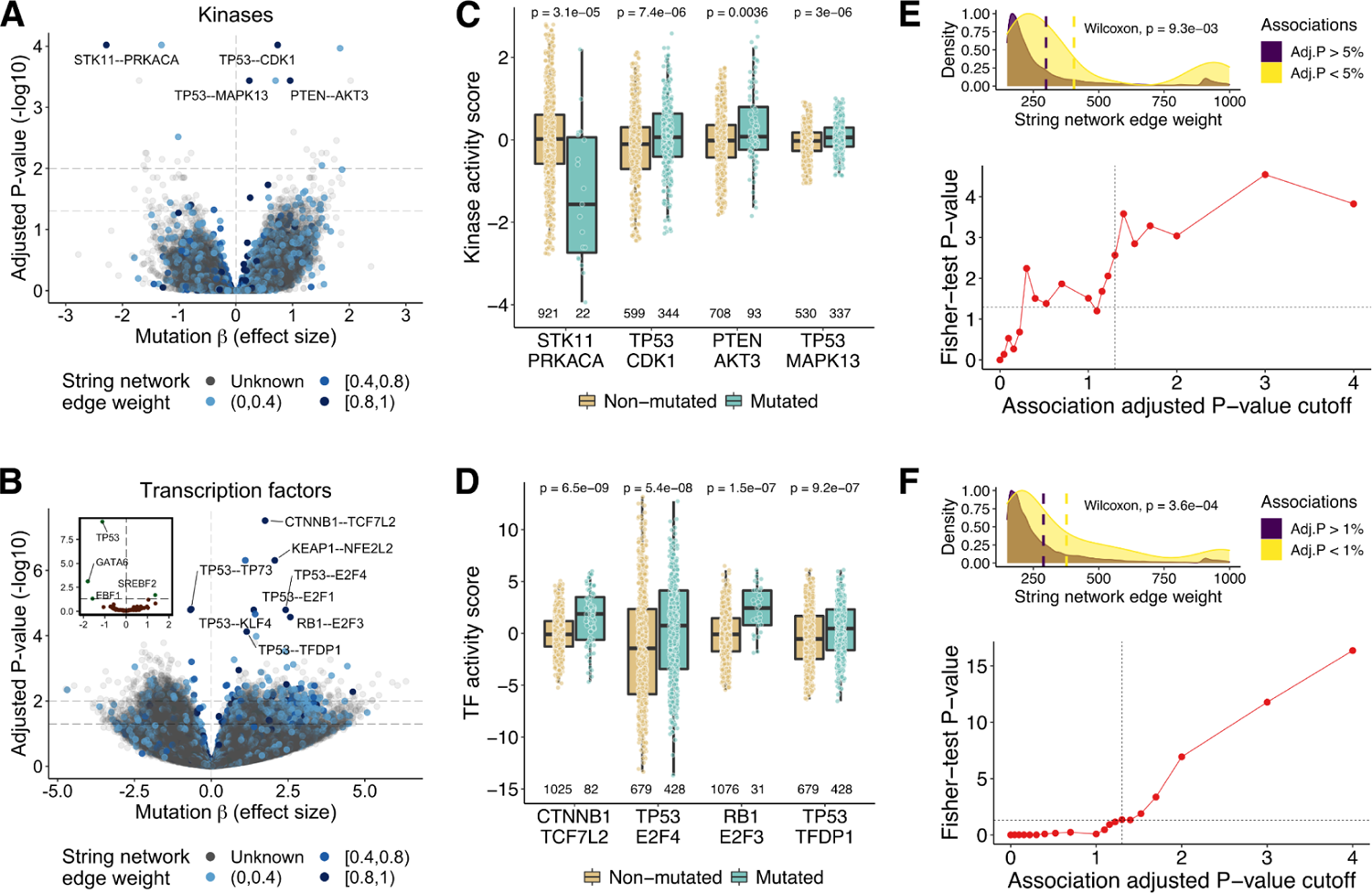
Genetic associations. **(A)** Volcano plot displaying the associations between the mutational status of genes and the activity of kinases. The x-axis contains the mutation coefficient (effect size) and the y-axis the adjusted P-values. The associations are represented in the form of a mutated gene - kinase. The color gradient represents the string network edge weight interval of the pair (grey if the pair is not in the string network). **(B)** Same as (A) for the TFs. The inner plot shows the effects of TF mutations on their own activities. **(C) (D)** Examples of the genetic associations highlighted in the volcano plots. The x-axis represents the associations and the y-axis the protein activities. The colors stratify the samples by their mutational status in the respective genes. The outliers (defined as the data points beyond Q1-1.5*IQR and Q3+1.5*IQR, where Q1 and Q3 are the first and third quartiles and IQR is the interquartile range) were removed from the distributions for representation purposes. The number of protein activity quantifications (including outliers) are shown beneath each boxplot. The P-values from Wilcoxon rank sum tests comparing both distributions are shown. All data points (including outliers) were used to calculate the P-values. **(E) Top panel.** Density plots comparing the edge weight distributions in the string network of the significant and non-significant association pairs obtained with the kinases. **(E) Bottom panel.** Enrichment of the associations in the string network (edge weight > 850) along multiple cutoffs of statistical significance. The x-axis shows the adjusted P-value cutoffs (-log10) and the y-axis the Fisher-test P-values (-log10). **(F)** Same as (E) for the TFs.

The association between mutated genes and altered protein activity contain several examples of previously known functional relationships. To evaluate this more broadly, we confirmed that our predicted associations were enriched in protein-protein functional associations annotated in the STRING database, both for the kinases and the TFs (P-value < 5%) (**Figures 2E, 2F - top plots**). We also performed an enrichment analysis using the string network along multiple cutoffs of adjusted P-values (**Methods**). The -log10 transformed P-values from the enrichment test increased as the association cutoffs were incremented (**Figures 2E, 2F - bottom plots)**, validating the generality of the significant associations. Overall, the genetic associations found are enriched in previously known functional associations, containing potential novel regulatory relationships for future experimental exploration.

### An atlas of kinase and TF regulation in cancer

The estimation of kinase and TF activities across a large set of tumour samples from different tissues provides a first look at the space of tumour signalling states as measured by hundreds of regulators. We projected the activity profiles in a lower-dimensional space using the uniform manifold approximation and projection for dimension reduction algorithm (UMAP) (**Methods**). For both the kinases and TF activities we observed that cancer samples were not clustered by experimental study (**Figures 3A, S7A**). The same was also observed using a principal component analysis (PCA) (**Figures S7B-S7C**). These results suggest that our normalization procedures helped to mitigate the technical biases between studies, being likely superimposed by biological variation.

**Figure 3.**
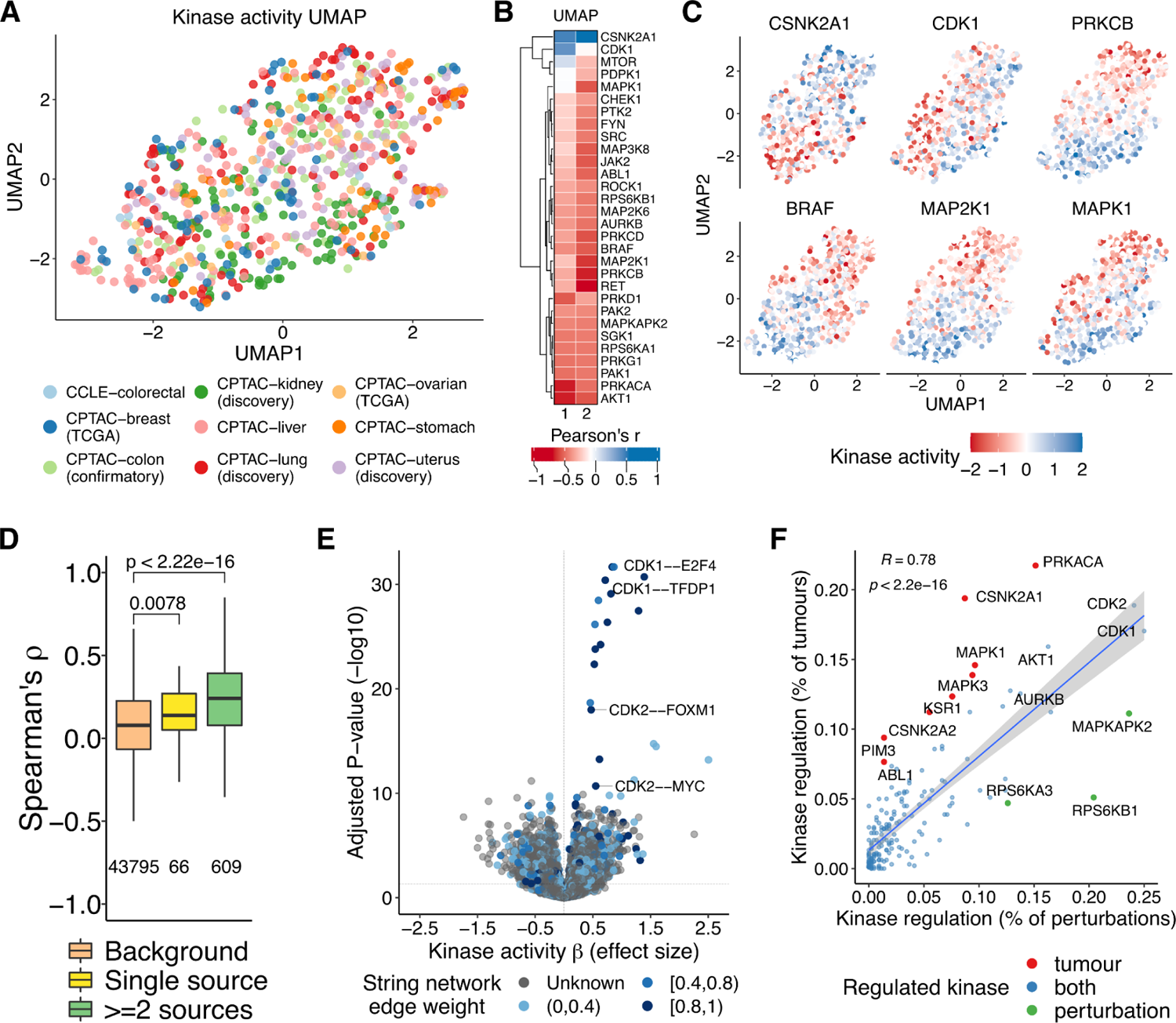
Regulation of protein activities in tumours and human perturbations. **(A)** UMAP projection of the kinase activity matrix (kinases as variables). The samples are colored by experimental study. **(B)** Pearson correlation coefficient between the UMAP projections and the activity of non-redundant highly variable kinases. **(C)** Kinase activity gradient along the samples for a selection of the kinases shown in (B). **(D)** Spearman’s rank correlation coefficients between the activities of kinases known to co-regulate each other. The pairwise kinase co-regulatory relationships were obtained from the OmniPath database and stratified by their presence in the OmniPath’s sources (as single source or in at least two different sources). We only kept activating and consensual interactions along the sources. The background corresponds to kinase pairs without known co-regulation events. The distributions were compared to the background using Wilcoxon rank sum tests. **(E)** Associations between the activity of kinases and TFs. The x-axis contains the kinase coefficients (effect sizes) and the y-axis the adjusted P-values. Each association is represented in the form of kinase - TF. The color gradient represents the edge weight of the pair in the string network (grey if not present). **(F)** Linear regression between the percentage of samples where the kinase is regulated in the perturbed conditions and in the tumour samples. The trend line and the Pearson’s r, with the respective P-value, are shown. In red and green are the kinases preferably regulated in the tumours and in the conditions, respectively. In blue are the kinases regulated in both.

After including only one kinase from sets of redundant kinases based on shared substrates such as AKT1/AKT2 (**Methods**), we selected the 30 kinases with the largest amount of variation along the samples (SD > median SD). As expected, these kinases are highly correlated with the UMAP projections (**Figures 3B**). This set of highly variable kinases contains known cancer drivers and kinases with inhibitors already used in the clinic as cancer treatment, such as BRAF, AKT, MAP2K1, SRC among others. Examining the tumour samples in this two dimensional representation indicates that highly regulated kinases in the same pathway tend to be activated or inhibited across the same samples (**Figure 3C)**. For example, we found that tightly co-regulated kinases from the MAPK signalling pathway, including PRKCB (PKC), BRAF (RafB), MAP2K1 (MEK1) and MAPK1 (ERK), share the same activation pattern along the cancer samples (**Figure 3C)**. CDK1 is known to phosphorylate the casein kinase 2 (CSNK2A1). These kinases together showed opposite correlations with the UMAP projections and, consequently, a distinct regulatory state across the samples (**Figures 3B-3C)**.

We obtained pairwise kinase regulatory relationships deposited in the OmniPath database (Türei, Korcsmáros, and Saez-Rodriguez 2016) and correlated their activities (**Methods**). We found that kinases that regulate each other were more likely to have correlated patterns of activity across samples (**Figure 3D**). This was still observed when taking into account cases where the pair of kinases shared some substrates (**Figure S7D; Methods**). Similarly, we would expect that kinases and TFs within the same pathway will tend to have similar patterns of activation across the samples. To investigate this, we modelled the TF activities as a function of the kinase activities using linear regressions (**Methods**), identifying 5,712 significant associations at an FDR < 5% (3,130 for FDR < 1%) (**Figure 3E; Table S4**). These associations were enriched in known kinase-TF functional interactions (**Figure S7E-S7F**), including for example the relation between CDK1 activity and the activities of E2F4 and TFDP1 (Spring et al. 2017; Jiao et al. 2017) (**Figure S7G**). Altogether, these results corroborate that the variation in activities across the samples is shaped to some extent by the underlying regulatory relationships.

Our analysis can indicate the kinases that are most often misregulated in cancer. For comparison, we also estimated kinase activity changes from phosphoproteomic measurements in a large panel of other conditions (Ochoa et al. 2016). We observed a correlation between the degree of regulation of kinases in cancer and non-cancer conditions (r=0.78, p-value=2.2e-16), with AKT1 and the cell-cycle kinases CDK1/2 and AURKB being highly regulated in both sets of conditions (**Figure 3F**). Kinases deviating from the regression line can be classified as preferentially regulated in the tumours or in the other conditions (**Methods**). There were a larger number of kinases specifically dysregulated in cancer (e.g including PRKACA, CSNK2A1 and MAPK1) compared with other non-cancer conditions (**Figure 3F**). The kinases MAPKAPK2, RPS6KB1 and RPS6KA3 were more often regulated in other conditions when compared with their degree of regulation in tumours (**Figure 3F**). We performed the same analysis by tissue type (**Figure S8A**). The number of specifically dysregulated kinases was consistently higher in the tumours than the non-cancer conditions in all tissues (**Figures S8A-S8B**). The inter-tissue variation regarding the number of dysregulated kinases in tumours correlated with the number of samples but not with the number of kinases quantified in the tissues (**Figure S8C**). Some kinases (e.g. PRKACA, CSNK2A1 and MAPK1) were found specifically dysregulated across multiple tumour types, but more than half were dysregulated in just one tissue (68%) such as MYLK kinase in stomach cancer and PIM3 in kidney cancer (**Figure S8D**).

Finally, using the activity profiles we clustered and stratified all samples into 8 cancer activity subtypes (**Methods**). We characterized each of the subtypes by performing over-representation analysis of clinical features and the activities that are most often regulated in each of the clusters. We then used CARNIVAL (Dugourd et al. 2021; A. Liu et al. 2019) to investigate the most plausible mechanistic links that could connect the most regulated kinase and TF activities in each cluster (**Methods**). We provide an extensive description of these 8 activity subtypes in **Supplementary Results**. Some of these activity subtypes are enriched in specific tissue or subtypes characterized by other approaches. For example, cluster 1 is enriched in high grade serous ovarian cystadenocarcinoma (SOC) with consistent activation of ARID1A. Cluster 5 is enriched in lung carcinoma and breast cancer samples with a general high activity of ZEB2, a promoter of epithelial to mesenchymal transition (EMT), metastasis and resistance in LUAD and breast cancer (Malvi et al. 2019; Bin Zhang, Zhang, and Shen 2015). As an interesting example, Cluster 7 was found to be enriched in CD8- inflammation tissues with consistent activation of the proinflammatory JUN and NFKB1 TFs likely via the increased activity of PAK1.

### Differential protein activity is associated with changes in patients survival

Survival analyses from multi-omics datasets have been largely based on mutation, gene or protein expression differences between groups of patients. However, kinase and TF activities should capture the signalling state of the cancer samples and could be also linked to overall patient survival (OS). To explore this we first performed a log-rank test to compare the Kaplan-Meier (KM) survival curves between patients with TF and kinase activities classified as inactive, neutral and active (**Methods**). We found several TFs and kinases significantly associated with OS in different tumour types (**Figures 4A-4B; Table S5**). For instance, the degree of MYC activity was correlated with OS in brain and liver cancers (**Figures 4C-4D**). In both cases, patients with high MYC activity showed less OS than patients with neutral and inactivated MYC (**Figures 4C-4D**). According to the literature, MYC overexpression is a poor prognosis factor in liver and pediatric brain tumours (Lin et al. 2010; Zheng, Cubero, and Nevzorova 2017). To take into account the effects of possible confounding covariates, we performed a multivariate Cox regression analysis using the protein activity scores as a predictor, while controlling for conventional clinical covariates and the genotype of recurrently mutated genes (**Methods**). Reassuringly, our findings with the log-rank tests were largely recapitulated with the Cox models (brain: hazard ratio (HR) = 1.50 (95% CI 1.27-1.76), adjusted P-value = 2.6e-5; liver HR = 1.17 (95% CI 1.06-1.31), adjusted P-value = 1e-2). Interestingly, we also found that high activity of FOXA1 and FOXM1 is a good and poor prognostic factor in liver cancer, respectively (FOXA1: HR = 0.69 (95% CI 0.55-0.85), adjusted P-value = 4.7e-3; FOXM1: HR = 1.39 (95% CI 1.17-1.66), adjusted P-value = 1.9e-3) (**Figures S9A-S9B**). These two proteins are known for their opposite role in hepatocarcinogenesis. On the one hand, elevated expression of FOXM1 promotes tumor cell proliferation and, on the other hand, FOXA1 inhibits tumour progression by suppression of PIK3R1 expression (He et al. 2017; Yu et al. 2016).

**Figure 4.**
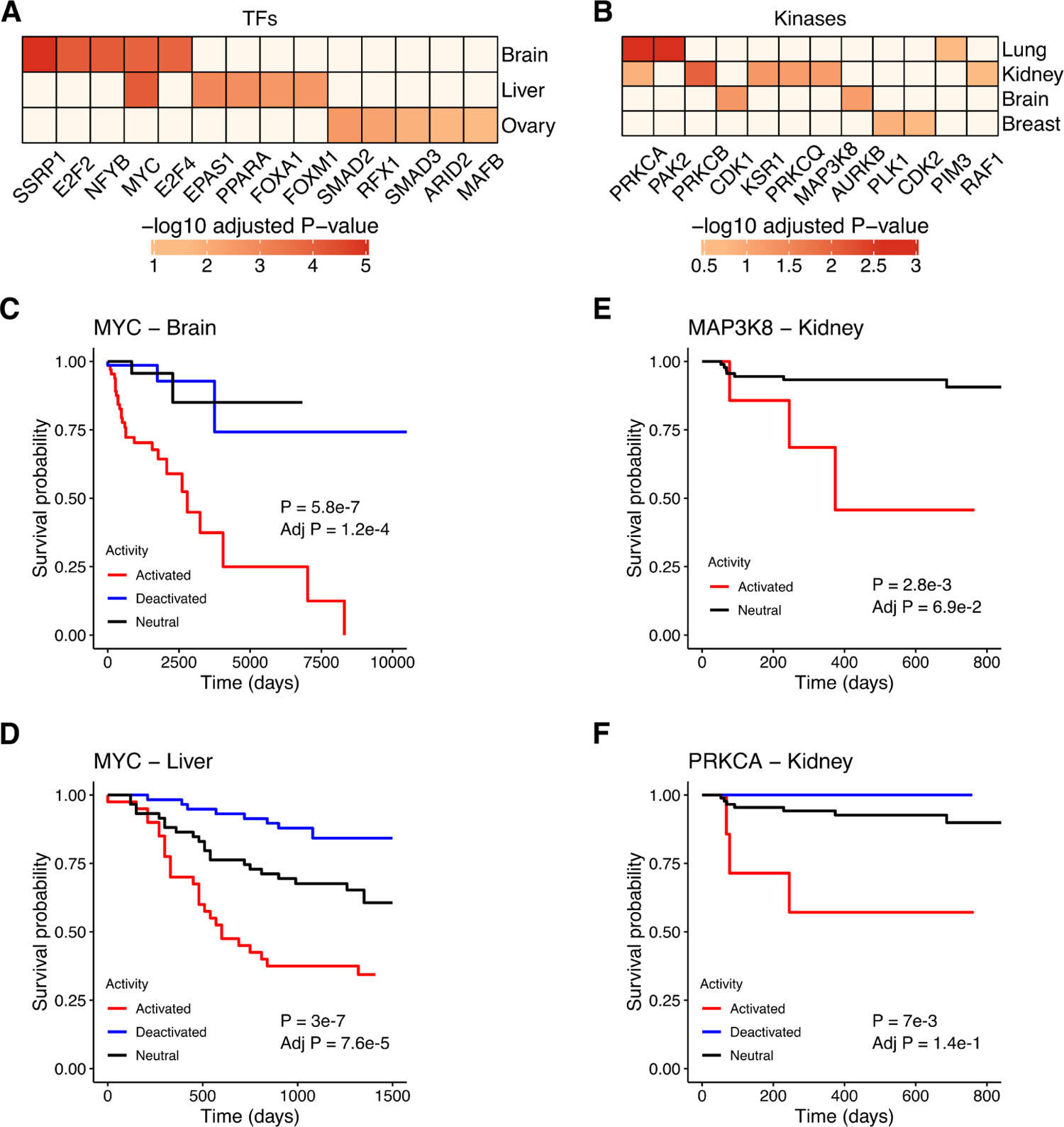
Survival analysis using kinase and TF activities. **(A)** Heatmap of log-rank test adjusted P-values (-log10) comparing Kaplan-Meier survival curves between cancer samples with TF activities classified as inactive, neutral and active (top 5 associations per tissue; FDR < 5%). **(B)** Same as (A) for the kinases (associations with FDR < 20%). KM survival plots comparing the survival probabilities (y-axes) as a function of time in days (x-axes) for MYC in **(C)** brain (inactive = 71, neutral = 39, active = 67) and **(D)** liver (58, 59, 40) cancer and **(E)** MAP3K8 (neutral = 95, active = 7) and **(F)** PRKCA (inactive = 3, neutral = 92, active = 7) in renal cancer. The log-rank P-values are shown in the plots.

Regarding the kinases, we found that elevated activity of MAP3K8 and PRKCA was associated with less probability of survival in renal cancer (MAP3K8: HR = 10.5 (95% CI 5.81-19), adjusted P-value = 1.1e-13; PRKCA: HR = 6.15 (95% CI 4.27-8.84), adjusted P-value = 2.3e-21) (**Figures 4E-4F**). In agreement with these results, the overexpression of both protein kinase C and mitogen-activated protein kinase 8 has been associated with a higher invasiveness of kidney tumours (Engers et al. 2000; J. Li and Gobe 2006; F. Liu et al. 2016; Su et al. 2015). Lastly, overactive AURKB was correlated with a lower survival rate in breast cancer (HR = 27.2 (95% CI 11.2-66.2), adjusted P-value = 5.1e-12) (**Figures S9C**), as previously found at the gene expression level (Huang et al. 2019). Altogether, these results indicate that the inference of kinase and TF activities can be a relevant prognostic tool in cancer studies.

## Discussion

Kinases and TFs are important mediators of cell signalling regulation and sensitivity to anti-cancer drugs. Here, we have compiled multi-omics datasets made available by the TCGA and CPTAC consortia and cell line studies and we were able to estimate the activities of 218 kinases and 292 TFs across 1,110 primary tumours and 77 cancer cell lines. Based on these we found that kinase activities appear to be primarily regulated by phosphorylation level with fewer cases of significant correlation with the predicted kinase protein abundance levels. Contrary to this, the predicted TF activity is primarily correlated with the mRNA/protein level of the TF itself with a smaller proportion of TFs with significant correlations with the phosphorylation state. This difference in regulation is not simply due to lack of detection of phosphosites as TFs have a median value of 6 phosphosites detected compared to 9 for kinases. A larger fraction of the TF activities is correlated with their mRNA levels than the protein abundance. This result is non-intuitive since the protein abundance should be a better proxy for activity. It is possible that this is due to the fact that TF activities are derived directly from the same mRNA datasets while there will be some degree of technical variation due to sample preparation and analysis when compared with the protein dataset.

Intuitively, the activity of a given protein (i.e., kinases and TFs) might be positively or negatively affected by mutations in the same protein or in other proteins it interacts with throughout the signalling networks. Our genetic analysis identified associations between mutated genes and the activities of kinases and TFs which were significantly enriched for known protein-protein interactions. Moreover, we found that the activities of the transcription factors TP53 and SREBF2 were correlated with their mutational status, as previously described (Garcia-Alonso et al. 2018). Nevertheless, we did not observe a general correlation between deleterious mutations within a kinase/TF and its activity including expected associations such as the BRAF V600E mutation and the activities of BRAF itself or other members of the MAPK pathway. These results were not due to issues linked with purity or immune infiltration as the same was observed in cell lines. Nevertheless there may be issues in linking mutations with signalling differences due to single-cell-level variances that can not be systematically profiled in bulk (Lun and Bodenmiller 2020). In the future, single-cell multi-omics profiling may allow us to consider intra-tumour genetic heterogeneities. We speculate that these observations are more likely explained by feedback loop mechanisms that are prevalent in signalling pathways (Lito, Rosen, and Solit 2013). These results emphasize the difficulty in interpreting the impact of mutations on signalling networks and the importance of studying directly the dysregulation of signalling in cancer (Yaffe 2019).

Known kinase regulatory pairs have strong patterns of co-regulation across the compiled dataset. These results suggest that the kinase activity estimates are meaningful, and that the variation in kinase activities along cancer samples is likely driven by biological factors. We showed in a previous work that these co-regulation signals can be used to predict kinase regulatory networks (Invergo et al. 2020). Similarly, we found many significant associations between the activities of kinases and TFs, which were significantly enriched in known functional interactions. This indicates that this compendium of protein activities may be useful in the future development of methods to reconstruct the signalling networks. Nevertheless, even the strongest correlations were modest in aggregate: well studied kinase-kinase regulatory pairs showed a median correlation of their predicted activities of 0.25 (**Figure 3D**). Our prior knowledge about kinase co-regulation is currently limited at multiple levels, including yet to be found regulatory relationships and the extent that these regulatory relationships depend on the tissue of origin or other factors. We speculate that these and other confounding factors could explain the weak kinase-kinase correlations.

By comparing the kinases most often differentially regulated across tumours and after more acute perturbations, we have shown that most often regulated kinases are the same in both contexts. These include kinases such as CDK1, AKT1 and AURKB. The most plausible explanation for this would be that kinases that are often regulated in acute perturbations are directly linked to the regulation of growth and cell-cycle and other critical processes needed to be regulated in cancer cells. Alternatively, we have shown that these highly regulated kinases occur in very central positions in the signalling network (Ochoa et al. 2016) and that it is possible that some degree of regulation of these kinases is almost unavoidable. Interestingly, this comparison allowed us to identify kinases which show higher differential regulation in tumours than acute perturbation such as PRKACA, MAPK1 and MAPK3.

Finally, we show how the estimated protein activity can be linked to differences in patient survival. Given that the activities of kinases and TFs can often be estimated via antibodies targeting regulatory phosphosites it may be possible to develop biomarkers based on these findings. In addition, kinases are very trackable drug targets with multiple kinase drugs already used to treat cancer patients. While further studies in cell based and animal models will be required to evaluate the significance of the findings presented here, this work provides kinase and TF activities linked to specific tumour types and mutational contexts that could be pursued for potential treatment.

## Methods

### Data collection

#### Proteomics and phosphoproteomics

The mass spectrometry (MS)-based protein and phosphosite quantifications (absolute [phospho]peptide intensities and ratios relative to controls) for the cancer samples of brain (Petralia et al. 2020), breast (Mertins et al. 2016), colorectal (Bing Zhang et al. 2014), kidney (Clark et al. 2019), liver (Gao et al. 2019), lung (Gillette et al. 2020), ovarian (H. Zhang et al. 2016), stomach (Mun et al. 2019) and uterus (Dou et al. 2020) were downloaded from the CPTAC data portal (proteomics.cancer.gov/data-portal). For the colon cancer samples (Vasaikar et al. 2019), we downloaded the data from the linkedomics database (linkedomics.org/login.php). The same data for the cancer cell lines of breast and colorectal tumours was downloaded from the respective publications (Lapek et al. 2017; Lawrence et al. 2015; Roumeliotis et al. 2017). The proteins and phosphosites were identified using gene symbols, in a process described by the common data analysis pipeline (CDAP) from CPTAC. Additionally, we downloaded normalized RPPA protein and phosphorylation quantification data (183 features across 7,694 samples from 31 TCGA tumours) from the TCPA (J. Li et al. 2013) database.

#### Transcriptomics

The RNA-seq data was obtained in the format of read counts and Fragments Per Kilobase of transcript per Million mapped reads (FPKM). The data for the tumour tissues of breast (Mertins et al. 2016), colorectal (Bing Zhang et al. 2014), kidney (Clark et al. 2019), lung (Gillette et al. 2020), ovarian (H. Zhang et al. 2016) and uterus (Dou et al. 2020) was downloaded from the GDC portal (portal.gdc.cancer.gov/). The data for the brain cancer was compiled from the pediatric cBioPortal (pedcbioportal.kidsfirstdrc.org/); for the liver (Gao et al. 2019) from NODE (www.biosino.org/node/) (accession ID: OEP000321); for the stomach (Mun et al. 2019) from GEO (ncbi.nlm.nih.gov/geo/) (accession ID: GSE122401); and for the colon cancer (Vasaikar et al. 2019) from the authors. The cancer cell lines (Lapek et al. 2017; Lawrence et al. 2015; Roumeliotis et al. 2017) data was downloaded from the CCLE data portal (portals.broadinstitute.org/ccle/data).

#### Genomics - somatic mutations

The whole genome sequencing (WGS)-derived somatic mutations for the brain cancer samples (Petralia et al. 2020) were downloaded from the pediatric cBioPortal (pedcbioportal.kidsfirstdrc.org/) in Mutation Annotation Format (MAF) files. For the breast (Mertins et al. 2016), colorectal (Bing Zhang et al. 2014) and ovarian (H. Zhang et al. 2016) cancers, the whole exome sequencing (WES)-derived MAF files were downloaded from the cBioPortal (cbioportal.org). The MAF file for the colon cancer samples (Vasaikar et al. 2019) was downloaded from the linkedomics database (linkedomics.org/login.php). Regarding the kidney (Clark et al. 2019), lung (Gillette et al. 2020) and uterus (Dou et al. 2020) cancers, we downloaded the MuTect2-called and VEP-annotated VCF files from the GDC data portal (portal.gdc.cancer.gov/). For the liver (Gao et al. 2019) and stomach cancers (Mun et al. 2019), we obtained the somatic mutations from the publication and authors, respectively. The mutation data for the colorectal and breast cancer cell lines (Lapek et al. 2017; Lawrence et al. 2015; Roumeliotis et al. 2017) was obtained from the DepMap portal (depmap.org/portal/).

#### Genomics - somatic copy number alterations

The somatic copy-number variation (CNV) data was downloaded as discretized GISTIC2 scores (Beroukhim et al. 2007; Mermel et al. 2011) and segment-level log2 ratios between the tumor and normal samples. The GISTIC2 scores can be −2 (strong copy-number loss, likely a homozygous deletion), −1 (shallow deletion, likely a heterozygous deletion), 0 (diploid), 1 (low-level gain of copy number, generally broad amplifications) and 2 (high-level increase in copy number, often focal amplifications). The GISTIC2 scores for the tumour samples of breast (Mertins et al. 2016), colorectal (Bing Zhang et al. 2014) and ovarian (H. Zhang et al. 2016), and for the cancer cell lines (Lapek et al. 2017; Lawrence et al. 2015; Roumeliotis et al. 2017) were downloaded from the cBioPortal (cbioportal.org). The same data for the brain cancer samples (Petralia et al. 2020) was downloaded from the pediatric cBioPortal (pedcbioportal.kidsfirstdrc.org/); for the colon samples (Vasaikar et al. 2019) from linkedomics (linkedomics.org/login.php); and for the liver samples (Gao et al. 2019) from NODE (biosino.org/node/index) (accession ID: OEP000321). The segment-level log2 ratios for the kidney (Clark et al. 2019) and uterus (Dou et al. 2020) cancer samples were provided by the authors of the respective publications.

#### Clinical data

The metadata and clinical information from the patients of breast (Mertins et al. 2016), colorectal (Bing Zhang et al. 2014) and ovarian (H. Zhang et al. 2016) cancers was obtained from the CPTAC (proteomics.cancer.gov/data-portal) and the cBioPortal (cbioportal.org) databases. The survival data for these patients was collected from (J. Liu et al. 2018), whereas the cancer subtypes were obtained from the respective publications and also using the *PanCancerAtlas_subtypes* function from the *TCGAbiolinks* R package (Colaprico et al. 2016). For the brain (Petralia et al. 2020), kidney (Clark et al. 2019), liver (Gao et al. 2019), lung (Gillette et al. 2020), stomach (Mun et al. 2019) and uterus (Dou et al. 2020) cancers, we downloaded the clinical information from the CPTAC portal and from the respective publications. For the colon cancer samples, we downloaded the data from the linkedomics database (inkedomics.org/login.php). The clinical data from the cancer cell lines donors (Lapek et al. 2017; Lawrence et al. 2015; Roumeliotis et al. 2017) was obtained from the CCLE data portal (portals.broadinstitute.org/ccle/data). Altogether, we collected the following information about the cancer patients: age, gender, ethnicity, race, height, weight, cancer histological type and subtype, tumour stage, overall survival and survival time in days.

### Data pre-processing and normalization

#### Proteomics

The label-free protein quantifications (precursor areas) for the colorectal tumours (Bing Zhang et al. 2014) and the tandem mass tag (TMT) protein intensities for the breast and colorectal cancer cell lines (Lapek et al. 2017; Lawrence et al. 2015; Roumeliotis et al. 2017) were pre-processed and transformed to log2 fold-changes as previously described (Sousa et al. 2019). For the brain (Petralia et al. 2020), lung (Gillette et al. 2020) and stomach (Mun et al. 2019) cancers, the sample replicates were combined by averaging the log2 fold-change values of each protein. After that, we removed 6 outlier samples from colorectal cancer with an absolute median log2 fold-change distribution higher than 1 (2-fold). Altogether, we assembled a matrix with 14,742 proteins and 1,266 samples (1,170 cancer samples and 96 cell lines) belonging to 9 different tissues. This matrix contained 9,941,918 protein measures (8,721,454 missing values) and 5,052 proteins quantified in at least 80% of the samples.

#### Phosphoproteomics

The phosphorylation measures were acquired at the phosphosite level. Each phosphosite is identified by a given protein, position and residue. The phosphosites from the different datasets were harmonized against a common reference by only keeping the phosphorylation sites that mapped correctly to the Ensembl human proteins (GRCh37 - release 98). As the phosphorylation sites were annotated at the gene symbol level (see data collection above), we mapped the phosphosites to the protein sequences using the canonical transcripts from UniProt (github.com/mskcc/vcf2maf/blob/main/data/isoform_overrides_uniprot). Duplicated phosphosites, arising from multiple phosphopeptide intensities mapping to the same phosphosite, were reduced to a single phosphosite if the log2 fold-change values were the same across all samples from the respective experimental study. All duplicated phosphosites were discarded otherwise. For the colorectal cancer cell lines (Roumeliotis et al. 2017), the relative TMT intensities (obtained by dividing the TMT intensities per the mean TMT intensity for each protein) were divided by 100 and transformed to log2. For the brain (Petralia et al. 2020), breast (Mertins et al. 2016), lung (Gillette et al. 2020) and stomach (Mun et al. 2019) cancers, the sample replicates were combined by averaging the log2 fold-change values of each phosphosite. We removed 52 outlier samples with an absolute median log2 fold-change distribution higher than 1 (2-fold). Then, the log2 fold-change distributions across samples were quantile normalized in order to ensure comparable distributions, using the *normalizeQuantiles* function from the *limma* R package (Ritchie et al. 2015). To detect phosphorylation changes that are independent of the protein abundance, we regressed-out the protein levels from the respective phosphosites using a multiple linear regression model. The phosphosite log2 fold-changes were set as the dependent variables while the protein log2 fold-changes, age and gender were set as the independent variables. The residuals from the linear model were the phosphorylation changes not driven by the protein abundance or other confounding effects (age and gender). The final phosphoproteomic matrix contained 86,044 phosphosites across 980 samples (930 cancer samples and 50 cell lines) from 9 different tissues. Due to the sparseness of the phosphorylation data (7,280,101 measures and 77,043,019 missing values), only 256 phosphosites were quantified in at least 80% of the samples (2,438 in 50%). For the downstream analyses we only considered the phosphosites (69,599) that were quantified from phosphopeptides phosphorylated at single positions.

#### Transcriptomics

The RNA-seq data (FPKMs and read counts) downloaded from the GDC and GEO websites (see data collection above) were converted to tabular formats using in-house R scripts. For the liver (Gao et al. 2019) and stomach (Mun et al. 2019) cancer samples, we calculated FPKM expression values from the RSEM (B. Li and Dewey 2011) expected counts using the *rpkm* function from the *edgeR* R package (Robinson, McCarthy, and Smyth 2010). We obtained the gene lengths by calculating the size (in base pairs) of the merged exons of each gene, using the *gtftools* python script (H.-D. Li 2018) (genemine.org/gtftools.php) and the GENCODE v19 human gene annotation (gencodegenes.org). After selecting the protein-coding genes, as described in the GENCODE v19 annotation, we removed the genes without expression (FPKM > 0) in at least 50% of the samples of the respective dataset. The FPKMs of each gene were subsequently log2 transformed (adding a pseudocount of 1 to avoid taking the log of 0) and converted to log2 fold-changes by subtracting the log2 median FPKM across samples. The log2 fold-changes were calculated for each dataset separately. The final gene expression matrix contained 17,056 genes across 1,187 samples and 9 tissues (1,110 cancer samples and 77 cell lines). 14,966 genes were expressed in at least 80% of the samples.

#### Genomics - somatic mutations

We processed the VEP-annotated VCF files from the kidney (Clark et al. 2019), lung (Gillette et al. 2020) and uterus (Dou et al. 2020) cancer samples using the *bcftools split-vep* plugin (samtools.github.io/bcftools/howtos/plugin.split-vep.html) with the following parameters: *-f ‘%CHROM\t%POS\t%REF\t%ALT\t%QUAL\t%FILTER\t%CSQ\n’ -d -A tab*. Only the mutations passing all quality filters (FILTER == “PASS”) were selected for downstream analyses. The mutations from these samples were then collected in a single text file using in-house bash scripts. In all datasets, we selected the mutations that were annotated using the canonical UniProt transcripts (github.com/mskcc/vcf2maf/blob/main/data/isoform_overrides_uniprot) and classified as frameshift and in frame insertions/deletions (Indels), missense, nonsense, stop codon loss (readthrough mutations) and splice site. All mutations (except splice site) were standardized against the Ensembl human proteins (GRCh37 - release 98) by filtering out those mutations whose reference (wild type) residues did not match the protein sequences in the mutation positions. The reference/mutated residues and protein position of the mutations were extracted from the HGVSp codes. In total, we collected 284,882 mutations in 17,305 protein-coding genes, across 1,168 samples (1,079 tumours and 89 cell lines) from 9 different tissues.

#### Genomics - somatic copy number alterations

GISTIC2 (version 2.0.23) was used to process the segment-level log2 ratios for the kidney (Clark et al. 2019) and uterus (Dou et al. 2020) cancer samples and define the gain/loss events of each gene (see data collection above), using the default parameter settings (*-genegistic* and *-savegene* parameters were both set to 1). After obtaining the discretized GISTIC2 CNV scores for each dataset, only protein-coding genes were selected as described in the GENCODE v19 human gene annotation. The final CNV matrix contained 16,520 genes across 1,025 samples and 7 tissues (947 tumours and 78 cell lines).

#### Normalization of gene and protein expression data

The confounding factor related to the experimental batch (e.g. CPTAC-breast, CPTAC-brain, etc.) was removed using a linear regression model. This model was implemented with the mRNA expression or protein abundance of a given gene as a dependent variable and the experimental batch as independent variable. The residuals from the linear model were the protein or mRNA variation not driven by the technical differences between cancer datasets.

### NCI60 and CRC65 cell lines - data collection and pre-processing

The proteomics and phosphoproteomics data for the NCI60 and CRC65 cancer cell lines (trypsin-digested version) were downloaded from (Frejno et al. 2020). The phosphorylation residues were obtained by mapping the position of the modifications to the UniProtKB/Swiss-Prot canonical and alternative (isoforms) protein sequences (release 2020_02) (uniprot.org/downloads). Only the phosphosites mapping to serine, threonine and tyrosine residues were selected. The log10 transformed phosphorylation and protein absolute abundances (iBAQ) were set to the original values using powers of 10 (10^*abundance*). Phospho(peptide) abundances mapping to the same phosphosite or protein were averaged per sample. The absolute abundances were converted to relative values (fold-changes) by calculating the log2 ratio of the abundances over the median abundance across cell lines. This process was performed for both cancer cell line sets. To detect net phosphorylation changes we regressed-out the protein levels from the respective phosphosites using the residuals of a linear regression model (y ∼ x) where the phosphosites were set as dependent variables (y) and the proteins as independent variables (x). In total, we assembled 11,940 proteins and 45,557 phosphosites across 125 cell lines (60 from NCI60 and 65 from CRC65).

The gene expression data was downloaded in the format of FPKMs (discover.nci.nih.gov/cellminer/) and Transcripts Per Million (TPMs) (depmap.org/portal/), for the NCI60 and CRC65 cell lines, respectively. Both gene expression measures were log2 transformed and converted to fold-changes by subtracting the log2 median FPKM/TPM across cell lines. In total, we calculated the log2 fold-changes of 18,291 genes across 95 cell lines (60 and 35 from the NCI60 and CRC65 sets, respectively). The whole genome mutation data was downloaded from the CellMiner (discover.nci.nih.gov/cellminer/) and the DepMap databases (depmap.org/portal/) for the NCI60 and CRC65 cancer cell lines, respectively. Across the NCI60 cell lines, we selected those mutations where more than 50% of the respective reads contained the alternative allele. In both datasets, we selected the mutations annotated as silent, missense, nonsense, stop codon loss, frameshift and in frame Indels. The protein position of the mutations and respective reference/mutated residues were obtained from the HGVSp codes. Altogether, we collected 585,904 mutations in 17,259 genes along 96 cell lines (60 from NCI60 and 36 from CRC65).

### Inference of kinase and TF activities

Kinase and TF activities were estimated using known kinase and TF regulatory targets. The kinase-substrate relationships were obtained from (i) ProtMapper (Bachman, Gyori, and Sorger 2019), a literature-based resource of kinase substrates annotated at the phosphosite level. The resource contains phosphorylation sites aggregated from five databases (BEL Large Corpus, NCI-PID, PhosphoSitePlus, Reactome and SIGNOR) and three text-mining tools (REACH, RLIMS-P and Sparser) and (ii) a collection of phosphosites derived from *in vivo (Hijazi et al. 2020)* and *in vitro* (Sugiyama, Imamura, and Ishihama 2019) experiments. Only the phosphorylation sites correctly mapped to the Ensembl human proteins (GRCh37 - release 98) were considered for the subsequent analyses. In total, we collected the phosphorylation targets of 573 kinases. The transcriptional targets of the TFs were compiled from the DoRothEA R package (v1.2.0), using only interactions annotated with confidence A, B and C. The kinase activities were inferred using a one sample z-test, which was shown to perform well (Hernandez-Armenta et al. 2017). The activity of a given kinase in a given sample was estimated as follows:

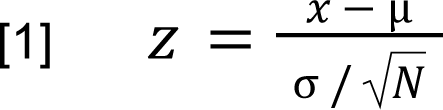

where *Z* corresponds to the z-score, *X* the average log2 fold-change of the kinase substrates, μ the average log2 fold-change of all phosphosites measured in the sample (background), √*N* the square root of the number of kinase substrates (*N*) and σ the standard deviation of the background. Then, the z-score was used to calculate a two-tailed P-value using the *pnorm* R function [2×pnorm(-abs(*Z*))], which was further log10 transformed and signed based on the position of the z-score in the standard normal distribution. If the z-score was in the right part of the distribution, i.e. positive, the kinase substrates showed an increase in phosphorylation in comparison to the sample background. Thus, the activity of that kinase was also expected to be increased (positive) in that sample, and vice-versa. This process was repeated for all kinases across all samples. For the downstream analyses we selected the kinase-substrate interactions from the databases and text-mining resources and the kinase activities quantified with 3 or more substrates, resulting in 218 kinases with activity estimates in an average of 437 cancer samples (of a total of 980 samples).

We also estimated kinases activities based on the phosphorylation changes of phosphosites mapping to the kinases. We selected the phosphosites with known regulatory status in PhosphositePlus or with unknown status but with a functional score higher than 0.4 (1,534 of 4,247 kinase-mapping phosphosites). The functional score was calculated using the *FunscoR* R package (evocellnet.github.io/funscoR/). The scores range from 0-1 and reflect the functional consequence of the phosphosites (Ochoa et al. 2020). The kinase activity inference method was the one sample z-test as described above. TF activities were estimated using the VIPER algorithm (Alvarez et al. 2016), using log2FC as gene level statistics (see Data pre-processing and normalization - transcriptomic). VIPER was run with a minimum limit of regulon size of 5, and using all provided gene level statistics as a background (eset.filter = FALSE). VIPER returned a normalized enrichment score for 292 TFs across 1,187 cancer samples.

### Benchmark of the kinase targets

We validated the kinase activities calculated from the different sources of kinase targets (database, text-mining, *in vivo* and *in vitro*) using a MS-based phosphoproteomic dataset reporting the relative phosphorylation changes of 52,814 phosphosites in 103 human perturbation-dependent conditions (Ochoa et al. 2016; Hernandez-Armenta et al. 2017). This data includes a gold standard dataset composed of 184 kinase-condition pairs where kinase regulation is expected to occur. The z-test-based absolute kinase activity scores estimated from the different kinase substrate sources were used as classifiers of kinase regulation. Given the imbalance between the positive (gold standard) and negative (kinase-condition pairs with unknown regulation) classes, we generated 100 random sets of negative cases with the size of the positive set. The predictive skill of each classifier was evaluated by the mean area under the receiver operating characteristic curves (AUROCs). As a control, we replicated the 100 random sets of negative and positive pairs (53 pairs each) along the different lists of kinase-substrates. The ROC curves and corresponding AUCs were calculated using the *prediction* and *performance* functions from the *ROCR* R package.

### Genetic associations with the kinase and TF activities

The effects of mutations on the kinase and TF activities were assessed by associating the activity of a given protein with the mutational status of the same protein or other proteins it might interact throughout the cellular regulatory networks. First, we built a binary mutation matrix M where the index M_ij_ corresponds to 1 if the sample i has a mutation in gene j and 0 otherwise. To do that, we selected the mutations classified as frameshift and in frame Indels, missense, nonsense and stop codon loss. Given the proteins X and Y, the association between the activity of Y (Y_act_) and the mutational status of X (X_mut_) was assessed across samples by fitting a linear model that took into account possible confounding effects:

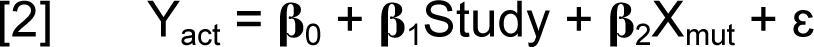

where Y_act_ represents the activity of protein Y, β_0_ the intercept, β_1_ the regression coefficient for the covariate experimental study, β_2_ the regression coefficient for the mutational status of X and ɛ the noise term. This model was applied to assess the effect of X_mut_ on the activity of the same protein (X_act_ ∼ X_mut_) and on the activity of other proteins (Y_act_ ∼ X_mut_). The P-values from the coefficients of X_mut_ (β_2_) were calculated using the *t*-statistic over a Student’s *t*-distribution and adjusted for false discovery rate (FDR) using the Benjamini-Hochberg method. The linear models and respective statistics were calculated using the *lm* and *p.adjust* R functions.

The associations were performed with the genes mutated in more than 20 samples and with the protein activities estimated in at least 10 samples. An association between a pair Y_act_ ∼ X_mut_ or X_act_ ∼ X_mut_ was performed if X_mut_ was mutated in at least 5 of all the samples in the pair. Regarding the Y_act_ ∼ X_mut_ associations, we tested 520,938 pairs between 208 kinases and 3,590 genes and 1,048,216 pairs between 292 TFs and 3,590 genes. In relation to the X_act_ ∼ X_mut_ associations, we tested 40 pairs and 64 pairs with the kinases and TFs, respectively.

### Projection of the protein activities in low-dimensional spaces

We reduced the dimensionality of the kinase and TF activity matrices using the PCA and UMAP methods. Given the sparseness of the kinase activity matrix, we imputed the missing values using the *missForest* function from the *missForest* R package. Prior to that, we selected the kinases (columns) with activity measures in at least 60% of the samples and the samples (rows) with measures in at least 80% of the kinases. The imputed kinase activity matrix contained 90 kinases across 727 samples. The PCA analysis was performed using the *prcomp* R function (scale. = T, center = T) and the UMAP analysis using the *umap* function from the *umap* R package (with default parameters).

When correlating the kinase activities with the UMAP projections (Pearson correlation coefficient), we excluded redundant kinases based on the degree of shared substrates. We first performed a hierarchical clustering analysis (*hclust* R function, agglomeration method = “complete”) using the Jaccard Index (JI) of shared substrates between kinases as distance measure (1-JI). Then, the kinase dendrogram was cut at a specific level (height = 0.85) to identify clusters of non-redundant kinases. We only kept one kinase per cluster (with the largest amount of substrates), reducing the number of kinases from 304 to 208.

### Correlation of kinase pairs

We obtained kinase-kinase regulation pairs from the OmniPath database (omnipathdb.org/interactions). We selected the interactions reported as directed, activating (stimulating relationships) and consensual along the resources (databases). Then, we correlated the activity of the kinase-kinase pairs along the samples using the Spearman’s rank correlation coefficient. The kinase pairs were stratified by the number of databases in which the interaction was found as a way of ascertaining the relevance of the interactions.

Strong correlations might be due to the amount of shared substrates between kinase pairs and not because of co-regulation events. To control for this technical limitation, we repeated the correlation analysis using a set of non-redundant kinases. This set was obtained by performing a hierarchical clustering analysis (*hclust* R function, agglomeration method = “complete”) using the degree of shared substrates between kinases as distance measure (1 - Jaccard Index of shared substrates). The dendrogram tree was cut at a height cutoff of 0.8. Just one kinase was kept per cluster (with the highest number of substrates). Using this approach, we reduced the number of kinases from 304 to 231.

### Associations between the activities of kinases and transcription factors

For a given protein pair K and T, where K is a kinase and T is a transcription factor, we tested whether the changes in the activity of kinase K are linearly associated with changes in the activity of the transcription factor T. To do that, we fitted a linear model to predict the activity of transcription factor T (T_act_) using the activity of kinase K (K_act_), while adjusting for possible confounding effects:

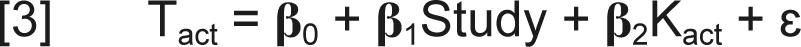

where T_act_ represents the activity of the transcription factor T, β_0_ the intercept, β_1_ the regression coefficient for the covariate experimental study, β_2_ the regression coefficient for the activity of kinase K and ɛ the noise term. The P-values from the coefficients of K_act_ (β_2_) were calculated using the *t*-statistic over a Student’s *t*-distribution and adjusted for false discovery rate (FDR) using the Benjamini-Hochberg method. The linear models and respective statistics were calculated using the *lm* and *p.adjust* R functions. Using this model we tested 26,280 kinase-TF associations between 90 kinases and 292 TFs.

### Enrichment of the protein association pairs in the STRING network

The genetic and the kinase-TF associations were tested for enrichment in the STRING protein-protein interactions network using Fisher’s exact tests (*fisher.test* R function, alternative = “greater”). The human network (version 11.0) was downloaded from the STRING database (string-db.org) as a list of protein-protein interactions with the corresponding combined scores. The scores range from 150 to 999 and represent the confidence of the respective interactions. We filtered the network using a minimum score of 850 to select the most confident protein-protein interactions. Fisher’s exact tests were performed by overlapping the protein association pairs with the STRING network across increasing -log10 adjusted P-values. The backgrounds corresponded to all the protein association pairs linearly modelled.

### Kinase activity changes between tumours and perturbations

To study the differences of kinase signalling between tumours and perturbation-dependent conditions, we estimated the activity of kinases across an extended panel of perturbations with phosphoproteomic measurements (Ochoa et al. 2016). This dataset is composed of 76,379 phosphosites across 439 perturbations. Next, we calculated the percentage of tumour samples and perturbations each kinase was regulated in, using an absolute kinase activity cutoff of 1.75 as previously used. We kept the kinases regulated in at least 1 tumour or perturbation. In order to find the kinases preferentially regulated in the tumours and in the perturbations - tumour or perturbation-specific kinases - we fitted a linear model between the percentage of kinase regulation in the tumours and in the perturbations, as independent and dependent variables, respectively. The most deviating kinases from the regression line were considered to be differentially regulated. These kinases were found by converting the residuals of the linear model to z-scores: while the kinases with a residual z-score > 2 were classified as tumour-specific, the kinases with a residual z-score < −2 were classified as perturbation-specific. This process was performed across all cancer samples and by tissue type. The linear models and respective residuals were calculated using the *lm* and *residuals* R functions. The residuals were standardized to z-scores using the *scale* R function.

### Survival analysis

In order to construct Kaplan-Meier (KM) survival curves, cancer samples were stratified based on their TF and kinase activity scores (AS). For each kinase and TF, we classified the samples as: inactive if AS < −1.75; active if AS > 1.75; neutral if −1.75 < AS < 1.75. The 1.75 activity cutoff was chosen based on a previous publication (Ochoa et al. 2016). We estimated the KM survival curves by protein and tissue. We tested if the differences on the activities of a given TF on a given tissue were associated with the probability of survival across time if: more than 10 deaths occurred and more than 10 samples were classified as active and inactive. Given the lower number of activation/inactivation events for the kinases, we tested the kinase-tissue pairs with more than 5 samples classified as active or inactive and with more than 5 deaths. These filters resulted in 1,025 tests for the TFs (274 TFs and 5 tissues) and 195 tests for the kinases (81 kinases and 7 tissues). The survival distributions of the cancer sample groups were compared using log-rank tests with the *survdiff* function from the *survival* R package. The P-values were adjusted for FDR using the Benjamini-Hochberg (BH) procedure (*p.adjust* R function). The KM curves were plotted using the *ggsurvplot* function from the *survminer* R package.

To account for confounding covariates, we performed a multivariate statistical analysis using Cox proportional-hazards regression models. The hazard function was fitted using the protein activity scores as a continuous predictor, adjusted for age, gender and the genotype (1 if mutated and 0 otherwise) of 28 recurrently mutated genes in our atlas (at least 100 mutations). Such models were applied to the protein-tissue pairs described above. We extracted the hazard ratios of the protein activity coefficients and corresponding 95% confidence intervals and P-values from the Cox models. The BH-corrected P-values were calculated using the *p.adjust* R function. The Cox regression models were fitted using the *coxph* function from the *survival* R package.

## Supporting information

Supplementary Results

## Acknowledgements

A.S. is funded by the FCT (Fundação para a Ciência e a Tecnologia) PhD Studentship PD/BD/128007/2016, under the GABBA program. A.D. was funded by German Federal Ministry of Education and Research (Bundesministerium für Bildung und Forschung BMBF) MSCoreSys research initiative research core SMART-CARE (031L0212A).

## Supplementary Figures

**Figure S1.**
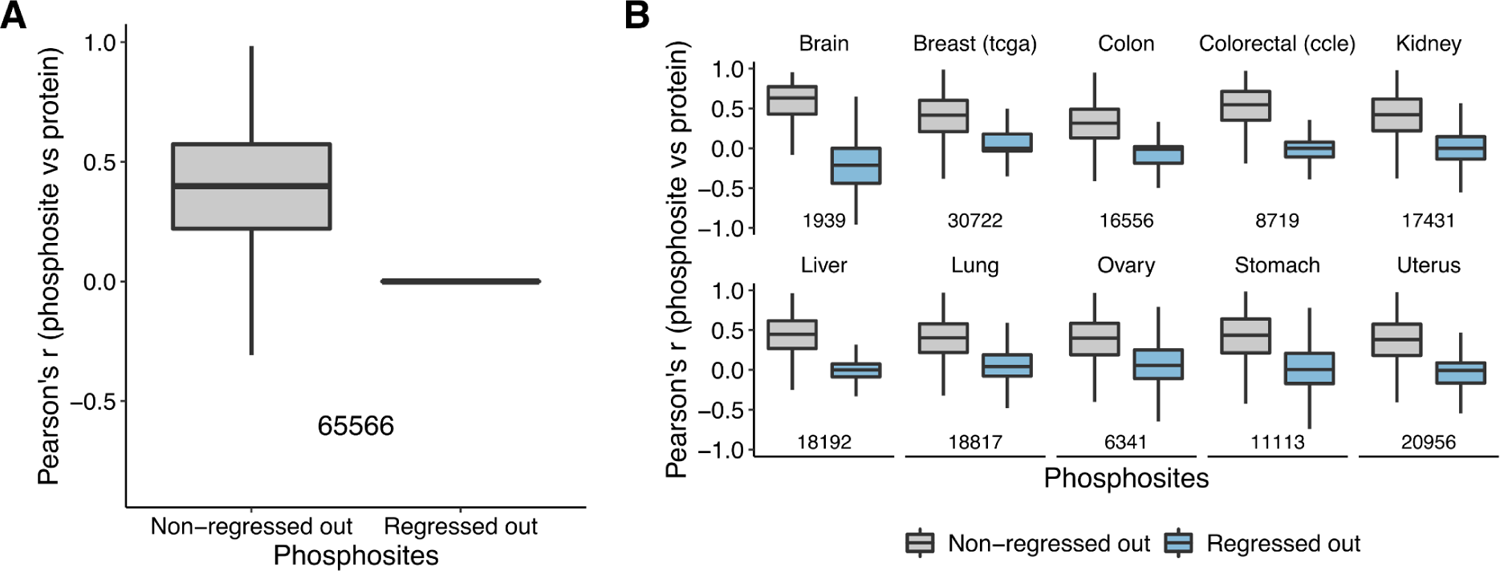
Pearson’s correlation between phosphorylation levels and corresponding protein abundances. **(A)** Distribution of the correlations between protein abundances and phosphorylation changes for protein-phosphosite pairs (number of pairs beneath the boxplots) across all cancer samples. Given the sparseness of the (phospho)proteomics data, we selected the protein-phosphosite pairs with protein/phosphorylation measures in at least 1% (n > 10) of the total cancer samples. Left: non-regressed-out phosphorylation data. Right: protein regressed-out phosphorylation data (**Methods**). **(B)** Representation of the same data as (A) by cancer dataset. Correlations were calculated for those protein-phosphosite pairs with protein/phosphorylation measures in at least 10% (n > 5) of the samples of each dataset. A small amount of correlation between phosphosites and proteins remain as the regression was done across all of the dataset.

**Figure S2.**
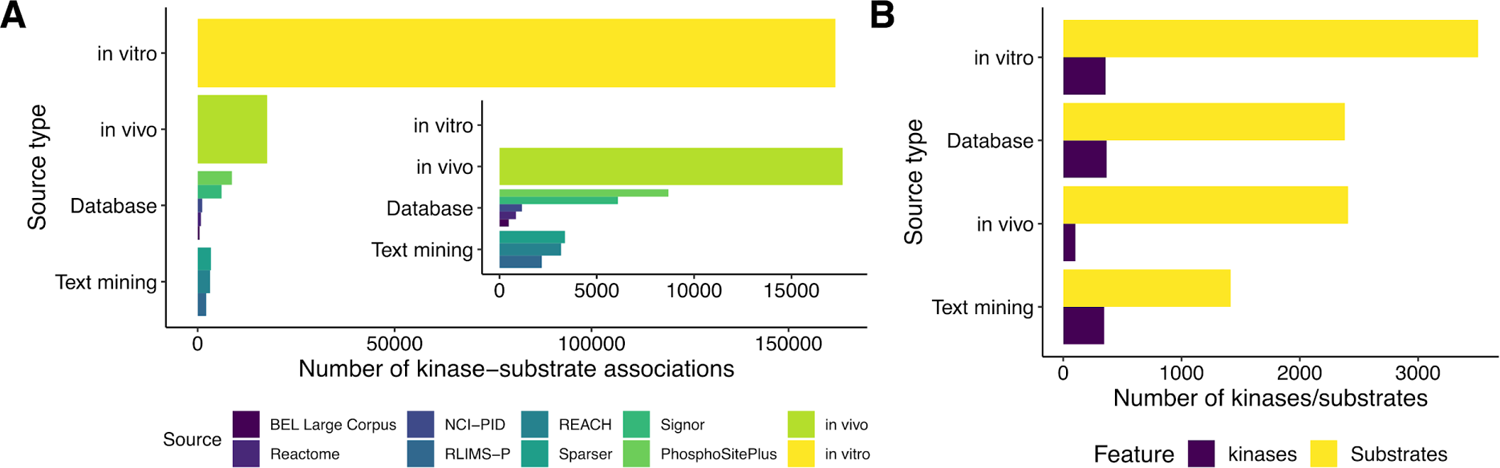
Lists of kinase-substrate associations compiled in this study. **(A)** Number of kinase-substrate associations by source type. **(B)** Number of kinases and substrates by source type.

**Figure S3.**
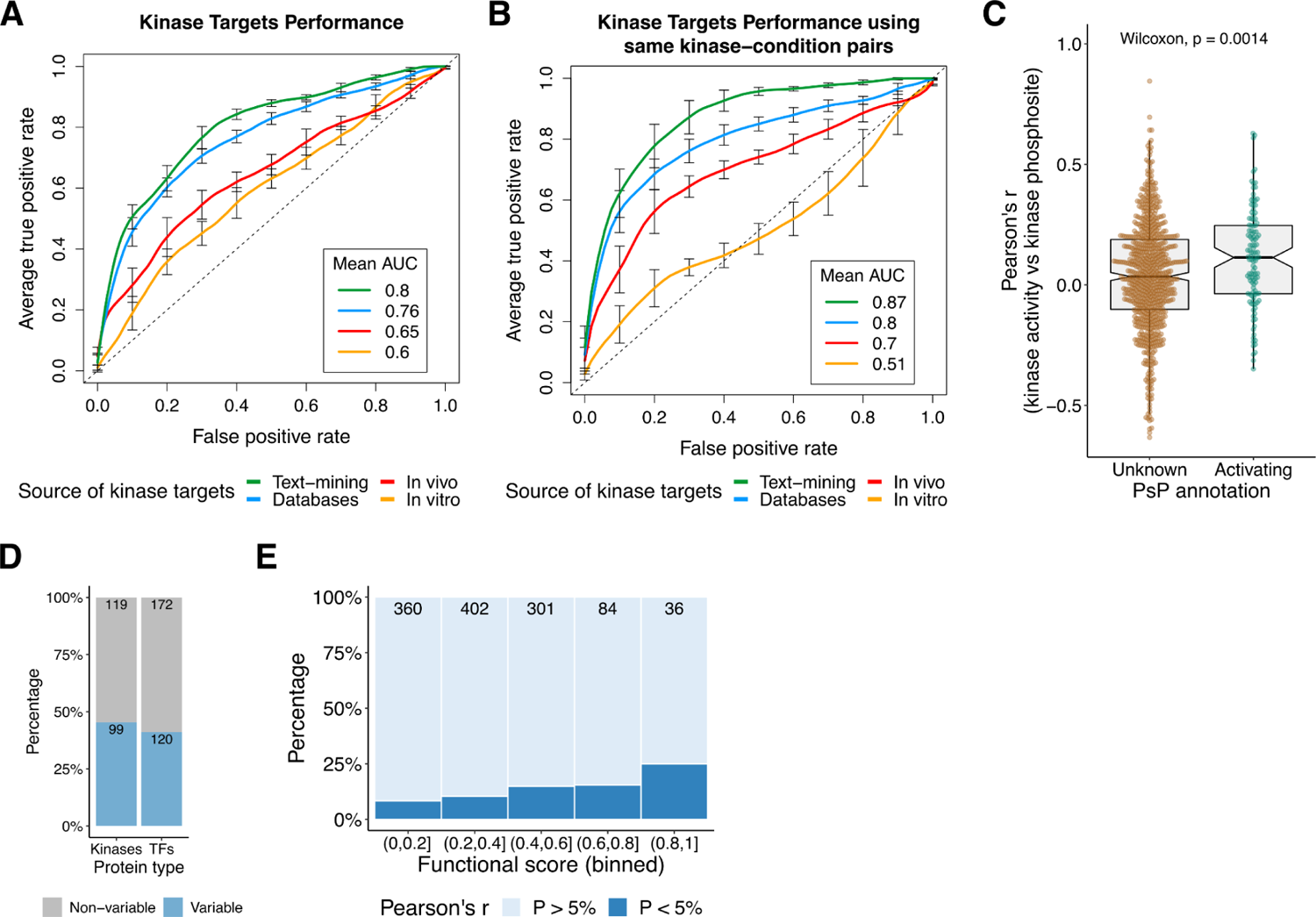
Validation of kinase-substrate sources and kinase activity estimates in the cancer samples. **(A)** Receiver operating characteristic (ROC) curves demonstrating the predictive performance of the Z-test-based kinase activities across different sources of kinase-substrate interactions. As positives, we used a set of 184 kinase-condition pairs where regulation is expected to occur, while as negatives we generated 100 random sets of the same size as the positive set. Curves display the average of 100 ROC curves and vertical bars the standard deviation of the true positive rate at multiple points of false positive rate. The average area under the ROC curve (AUC) is shown for each kinase-substrate list. The averaged ROC curves and corresponding AUCs demonstrate the discriminative power of each kinase-substrate list. **(B)** In contrast to the analysis shown in (A), here we replicated the 100 sets of negative (53) and positive (53) regulatory pairs along the different lists of kinase substrates. **(C)** Related to the main Figure 1C. Kinase activities were re-estimated in cancer samples after removing the kinase auto-regulatory phosphosites from the kinase targets. The boxplots show the distribution of the Pearson’s correlation between kinase activities and phosphosite quantifications that mapped to the same kinase, with (n = 118) and without (n = 743) annotation (activating) in PhosphoSitePlus. **(D)** Fraction of kinases and TFs classified as highly variable across the tumour samples. Kinases and TFs with absolute activity measures higher than 1.75 and 3.89 (96.7^th^ percentiles), respectively, in at least 5% of the samples were classified as highly variable. **(E)** Percentage of phosphosites in TFs significantly and not significantly correlated with the corresponding TF activities, stratified by their functional score. Analysis based on 1,183 phosphosites mapping to 178 TFs (n > 10).

**Figure S4.**
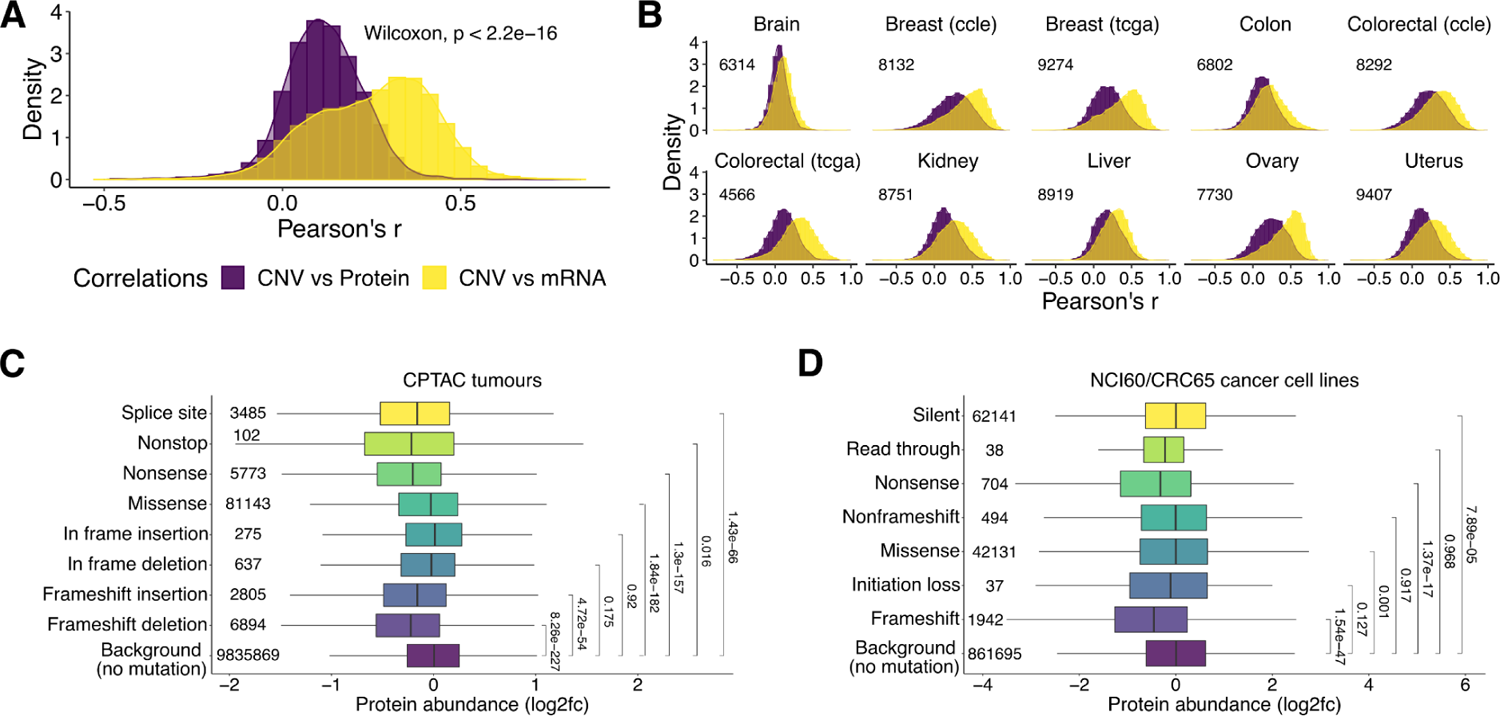
Effects of genomic alterations on protein abundances. **(A)** Comparison of the distribution of the correlations (Pearson’s r) between the CNV levels (GISTIC2) and the mRNA and protein abundances (log2 fold-changes). Correlations were calculated for those genes with CNV, mRNA and protein quantifications in at least 10 samples (11,624 genes). The experimental batch was regressed-out from the mRNA and protein quantification data before computing the correlations (**Methods**). **(B)** Same as (A) by tissue and experimental study. P-values < 2.2e-16 in all cases (Wilcoxon rank sum test). The number of genes is indicated in the plot. **(C)** Protein abundance distribution between mutation types from the CPTAC tumours. One sample may have multiple mutations in the same protein. Therefore, we selected the sample-protein pairs that were exclusive of each mutation type to prevent the cases where different mutations in the same protein and sample have the same protein abundance. The outliers (defined as the data points beyond Q1-1.5*IQR and Q3+1.5*IQR, where Q1 and Q3 are the first and third quartiles and IQR is the interquartile range) were removed from the distributions for representation purposes. The number of protein quantifications (including outliers) is shown at the left of each boxplot. The P-values from a two-sample T-test comparing each distribution with the background (no mutation) are shown at the right. All data points (including outliers) were used to calculate the P-values. **(D)** Same as (C) for the cancer cell lines from the NCI60 and CRC65 panels.

**Figure S5.**
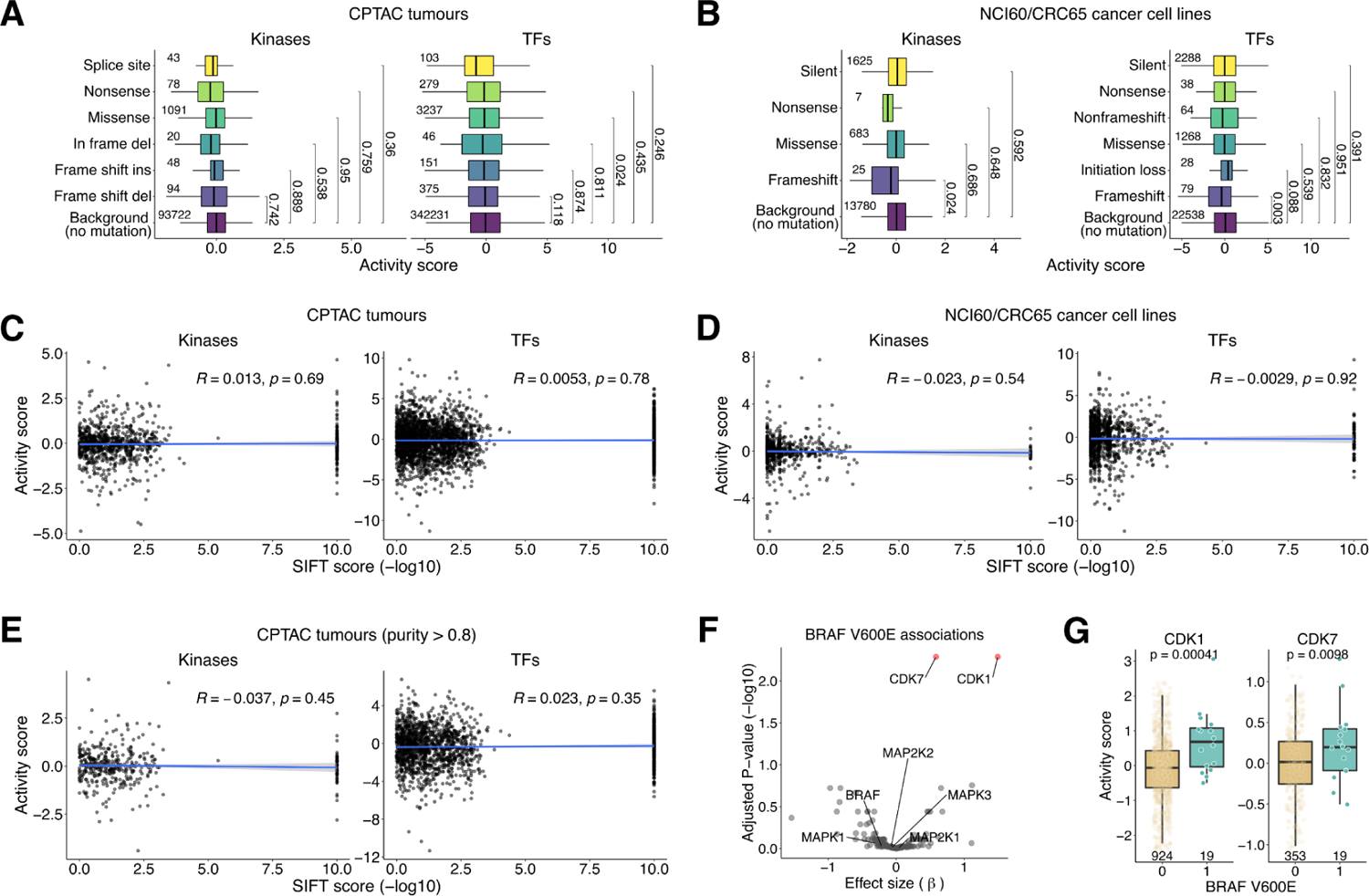
Effects of genomic alterations on protein activities. **(A)** Distribution of kinase and TF activities between mutation types from the CPTAC tumours. Only the sample-protein pairs that were specific of each mutation type were selected to prevent the cases where different mutations in the same protein and sample have the same protein activity. The outliers (defined as the data points beyond Q1-1.5*IQR and Q3+1.5*IQR, where Q1 and Q3 are the first and third quartiles and IQR is the interquartile range) were removed from the distributions for representation purposes. The number of protein activity quantifications (including outliers) is shown at the left of each boxplot. The P-values from a two-sample T-test comparing each distribution with the background (no mutation) are shown at the right. All data points (including outliers) were used to calculate the P-values. **(B)** Same as (A) for the cancer cell lines from the NCI60 and CRC65 panels. **(C)** Scatterplots between the -log10 SIFT score (x-axis) of missense mutations and the activity of kinases and TFs (y-axis) from the CPTAC tumours. The linear regression line and the Pearson correlation coefficient, with the respective P-value, are shown. Cases where the same sample had multiple missense mutations in the same gene were removed to prevent the assignment of the same protein activity to different SIFT scores. **(D)** Same as (C) for the cancer cell lines from the NCI60 and CRC65 panels. **(E)** Same as (C) for the CPTAC tumours with higher purity (greater than 0.8). The purity score provides information about the degree of immune infiltration and was calculated from the gene expression data using the ESTIMATE algorithm. **(F)** Volcano plot showing the associations between the BRAF^V600E^ mutation and the activity of kinases. The x-axis contains the mutation coefficient (effect size) and the y-axis the adjusted P-values. Highlighted are kinases from the MAPK/ERK signaling pathway and CDK1/7 (significantly associated with the BRAF^V600E^ mutation). **(G)** Differential activity of the CDK1 and CDK7 kinases between samples with and without BRAF^V600E^ mutation. The x-axis separates the cancer samples by mutation status (1 if mutated and 0 otherwise) and the y-axis contains the kinase activities. The outliers were removed from the distributions for representation purposes. The number of quantifications (including outliers) are shown beneath each boxplot. A P-value from a Wilcoxon rank sum test comparing both distributions is shown. All data points (including outliers) were used to calculate the P-values.

**Figure S6.**
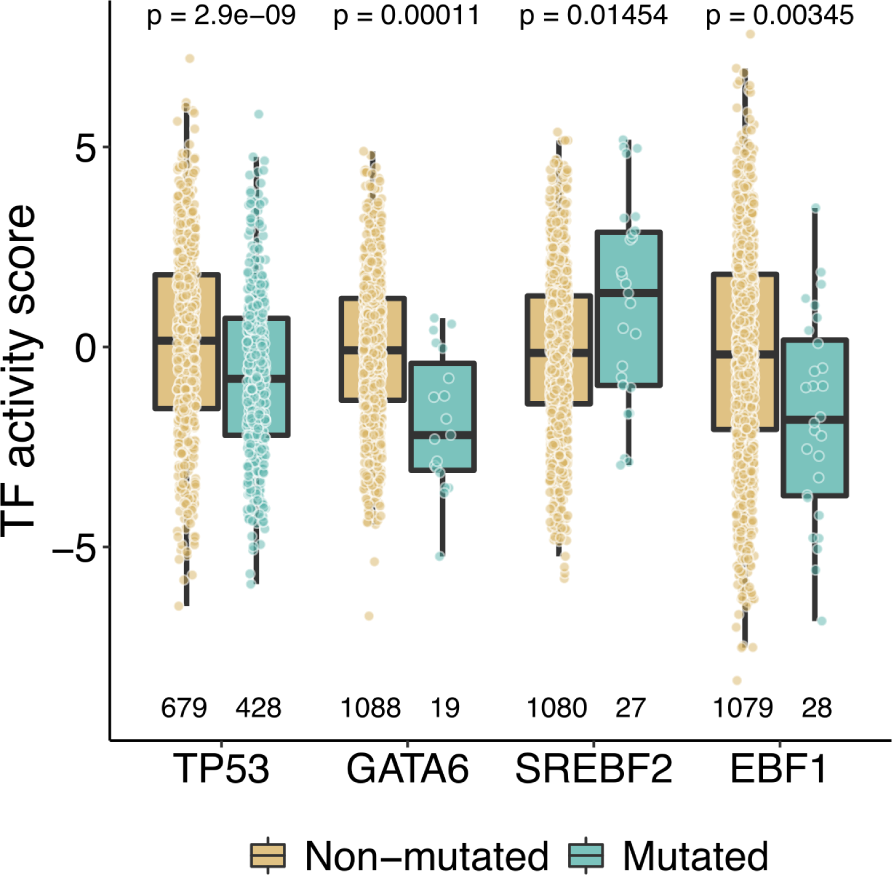
Examples of associations between the mutational status of TFs and their activities. Related to the main Figure 2B. The x-axis represents the TFs and the y-axis the activities. The colors stratify the samples by their mutational status in the respective TFs. The number of quantifications are shown beneath each boxplot. The P-values from Wilcoxon rank sum tests comparing both distributions are shown.

**Figure S7.**
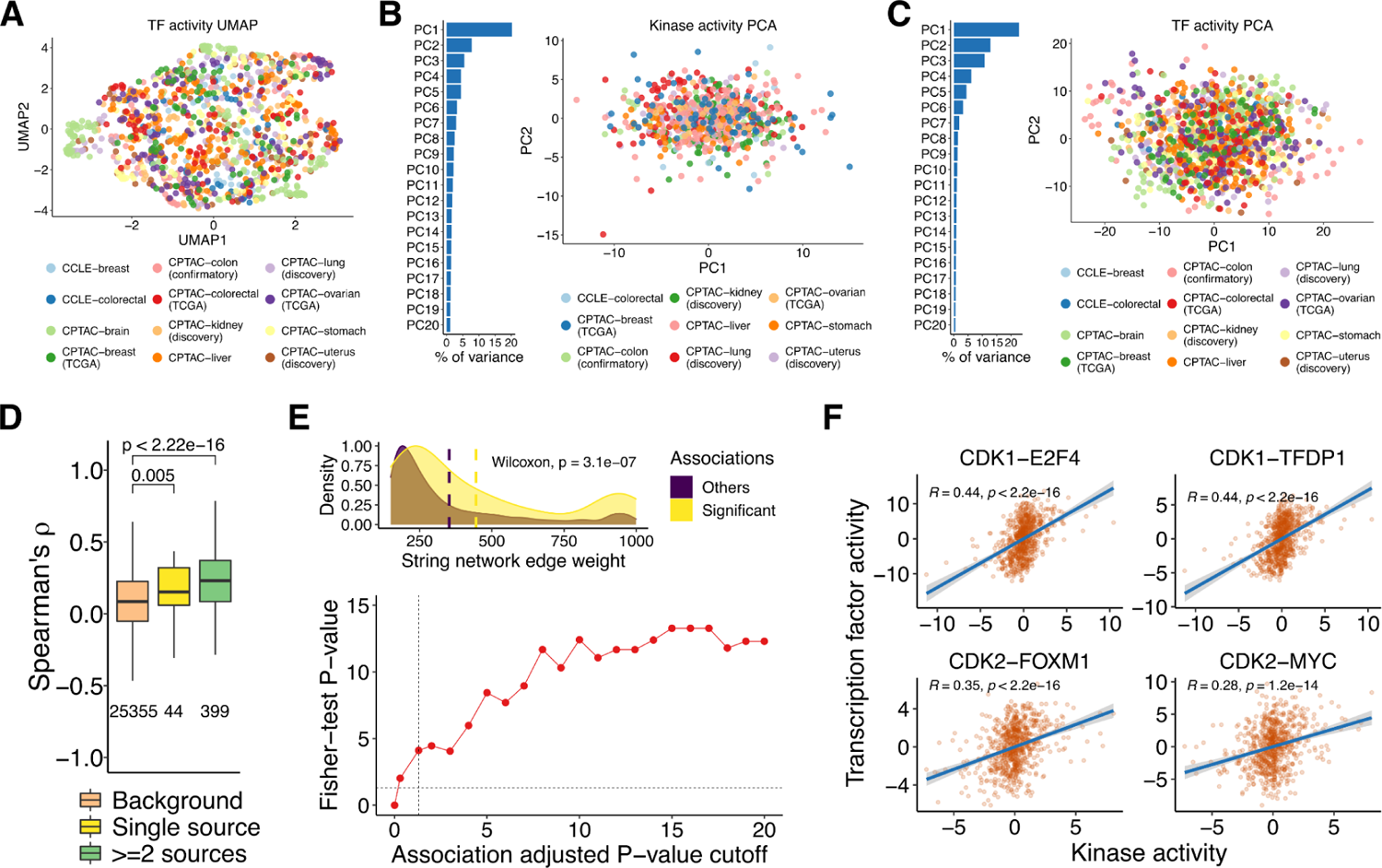
Projection of protein activities in low-dimensional spaces and kinase-TF associations. **(A)** UMAP projection of the TF activity matrix (TFs as variables). The samples are colored by experimental study. **(B)** PCA of the kinase activities. The barplots indicate the percentage of total variance explained by the first 20 principal components (PCs) (out of 90 PCs). The scatter plots illustrate the samples projected along the PC1 and PC2. The samples are colored by experimental study. **(C)** Same as (B) for the TFs. The barplot contains 20 of 292 PCs. **(D)** Related to the main Figure 3D. Correlations between the activities of non-redundant kinases with co-regulatory relationships. The co-regulatory interactions were obtained from OmniPath (activating and consensual interactions along the sources) and catalogued as present in a single source or in at least two different sources. The background corresponds to kinase pairs for which co-regulation is not known. The distributions were compared to the background using Wilcoxon rank sum tests. **(E) Top panel.** String network edge weight distributions between the significant and non-significant kinase-TF associations (224 and 7527 pairs). The significant associations were selected with a FDR < 5% and an absolute effect size > 0.5. **(E) Bottom panel.** Enrichment of the kinase-TF associations in the string network (edge weights > 850). The y-axis shows the Fisher-test P-values (-log10) and the x-axis the adjusted P-value cutoffs (-log10) that were used to select the associations. **(F)** Scatter plots of the kinase-TF associations highlighted in the main Figure 3E.

**Figure S8.**
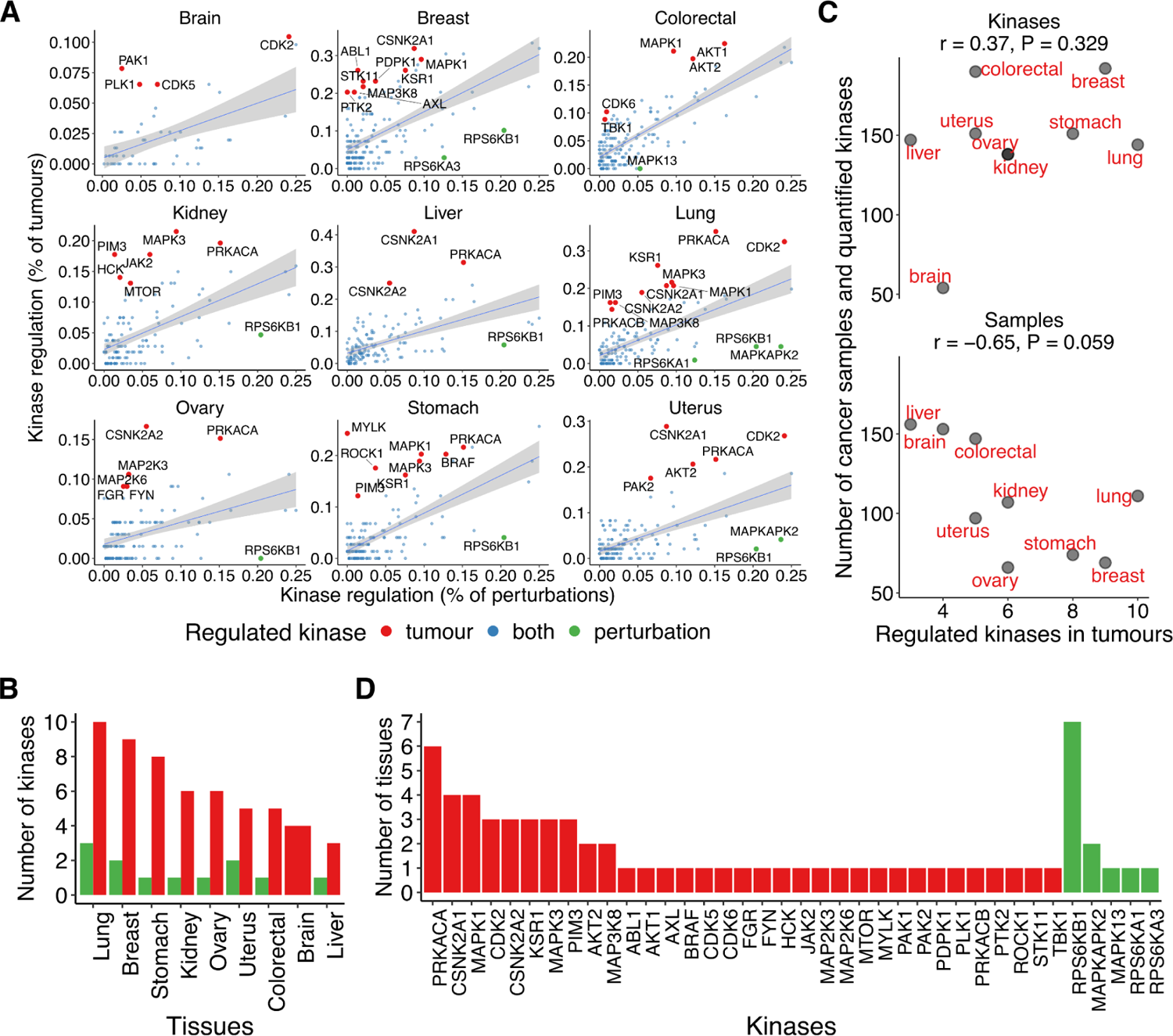
Kinase activity regulation in tumours and perturbed human conditions. **(A)** Related to the main Figure 3F. Linear regression models between the percentage of kinase regulation in the perturbed conditions (x-axis) and in the tumour samples (y-axis) by tissue type. **(B)** Number of kinases classified as regulated in the tumours (red) and in the conditions (green) in each tissue. **(C)** Correlation between the number of regulated kinases in tumours (x-axis) and the number of quantified kinases and samples (y-axis) across tissues. The Pearson’s r and respective P-value are shown. **(D)** Number of tissues where the kinases were identified as regulated in the tumours (red) or in the conditions (green). The kinases are mutually exclusive between them (no kinase found as regulated in the tumours and in the conditions).

**Figure S9.**
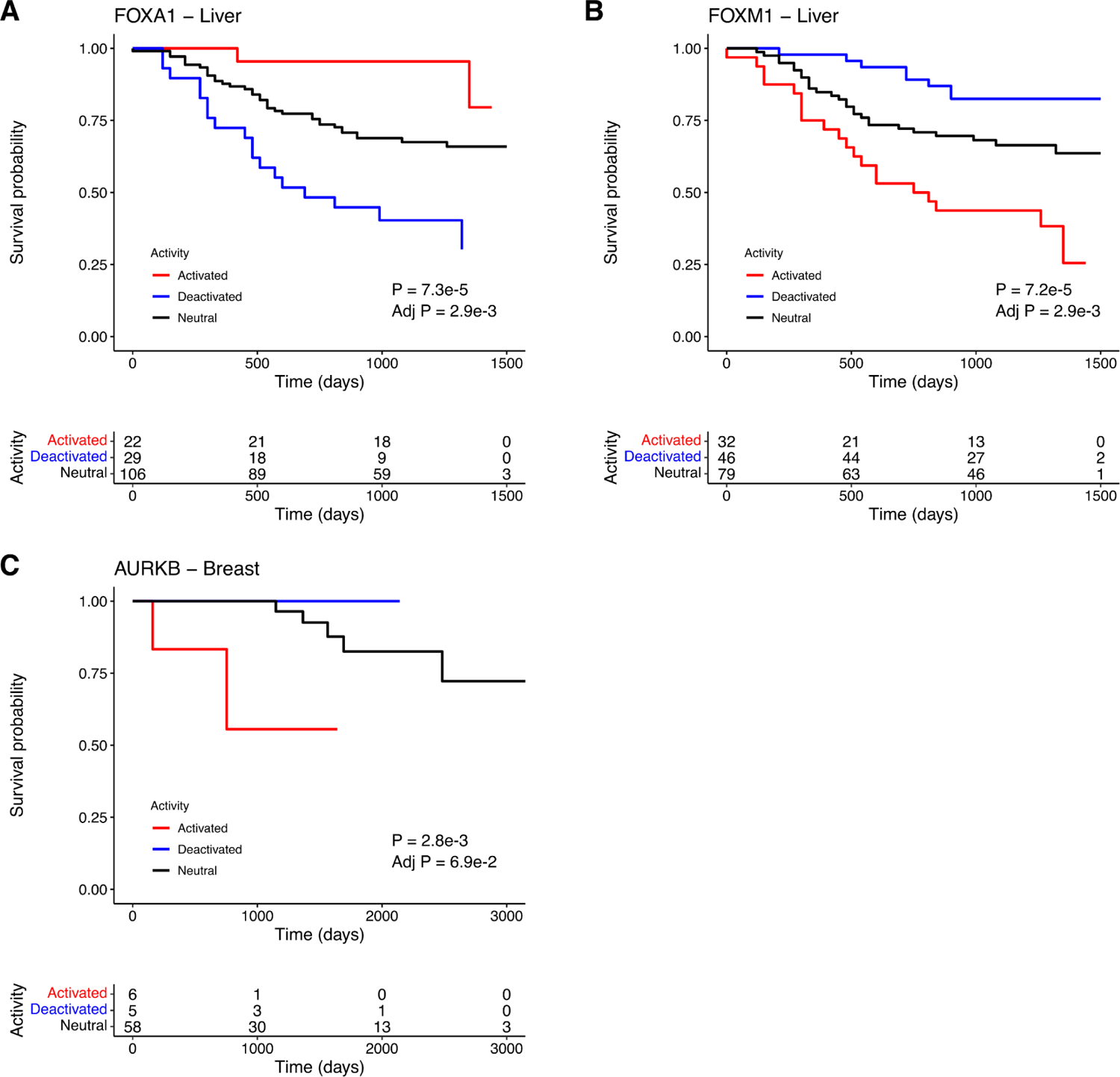
The activities of FOXA1/FOXM1 and AURKB are associated with the overall survival of liver and breast cancer patients. Related to the main **Figures 4A-4B**. KM survival plots for **(A)** FOXA1 (inactive = 29, neutral = 106, active = 22) and **(B)** FOXM1 (46, 79, 32) in liver cancer and **(C)** AURKB (5, 58, 6) in breast cancer. The tables beneath each plot contain the number of individuals at risk across time. The log-rank P-values are shown in the plots.

## Supplementary Tables

Table 1. Kinase and TF activities estimated from the molecular data.

Table 2. Significant correlations between the protein activities and the corresponding CNV, RNA, protein and phosphorylation levels.

Table 3. Genetic associations with protein activities.

Table 4. Associations between kinase and TF activities.

Table 5. Protein activities significantly associated with the overall survival of the cancer patients.

