## Supplementary Results for "Pan-Cancer landscape of protein activities identifies drivers of signalling dysregulation and patient survival"

**Supplementary results - Multi-omic based stratification of tumor samples.**

We clustered tumor samples according to kinase and transcription factor activities. To do so, we built a cross correlation matrix of tumor samples (by correlating their combined signature of batch corrected kinase and TF normalised enrichment scores (NES)) (Supplementary Results Fig 1). Based on visual inspection of the cross-correlation matrix, as well as information from applying silhouette and "within sum of square" methods, we decided to stratify tumor samples into 8 distinct groups (see methods).

To characterise the clusters, we performed an over-representation analysis of clinical features in each cluster (Supplementary Results Fig 2), to identify them with interpretable phenotypic information. This can help not only classify patients' clinical features based on their corresponding kinase and TF activity profiles, but also find functional similarities between samples that have different annotations. For example, The first cluster seems to be over-represented with ovarian cancer samples, thus other samples that are not ovarian cancers but where found in cluster 1 may still have relevant functional features that resemble ovarian cancer. Cluster 1 seems to be further over-represented with proliferative and stage iiic cancer types.

Cluster 2 seems to be over-represented with stage i lung cancer samples. Cluster 3 seems to be especially over-represented both serous/mucinous ovary and colon cancers, while a specific enrichment in colon cancer is found for cluster 4. Cluster 5 seems to be over-represented with breast cancer samples. Cluster 6 is marginally over-represented in kidney cancer samples, and CD8+ inflamed samples. Cluster 7 seems to be over-represented with inflammed (CD8-) mesenchymal and infiltrating lobular carcinoma, but no specific organ is found over-represented. Cluster 8 seems to be over-represented with basal-like, Uterine Corpus Endometrial Carcinoma with high CNV.

**TF and kinase activity characterization of tumor sample clusters**

In order to clearly characterise which enzyme and TF are the most consistently deregulated in each cluster, the mean of each kinase was divided by its corresponding standard deviation, within each cluster respectively (Supplementary Results Fig 3). Cluster 1 and 3 (over-represented with high grade serous ovarian cystadenocarcinoma (SOC)) both show a consistent activation of ARID1A. ARID1A is very frequently mutated with loss of function (but not loss of protein expression) in ovary cancers, except in high grade serous cystadenocarcinoma (Yachida et al. 2020). IRF1 down-regulation in cluster 1 is coherent with finding that its over-expression in SOC is associated with better prognosis and survival (Cohen et al. 2014). Additionally, NFKB has been found to have a tumor suppressor role in low grade SOC, while we observe a coherent down-regulation of NFKB1, RELA and SPI1 in cluster 1. Cluster 3 (also over-represented in high grade SOC), displays a different TF and kinase dysregulation profile (except for ARID1A). Cluster 3 shows an up-regulation of KDM5B activity, previously found to be associated with high grade SOC and poor prognosis (L. Wang et al. 2015). TF4AP is a transcription factor associated with SOC proliferation activated down-stream of oestrogen signaling, and is also up-regulated in cluster 3 (O'Donnell et al. 2005).

In cluster 2 (lung adenocarcinoma (LUAD) over-representation), BHLHE40 is consistently up-regulated, coherently with its recent reports of the up-regualtion of its activity in lung and esophageal carcinoma (Kiss, Mudryj, and Ghosh 2020). E2F6, which transcript levels are often elevated in LUAD, was also found to be up-regulated. Interestingly, MAZ and MTA2 are down-regulated in cluster 2, even though they are suspected drivers of metastasis in LUAD (Malvi et al. 2019; Zhang, Zhang, and Shen 2015). This could be explained by the fact that cluster 2 is actually over-represented with low grade (stage i and ia) tumors. Cluster 5 is also over-represented with LUAD, but also with breast cancer (BRCA). It shows a very different profile of TF/kinase dysregulation than cluster 2. Cluster 5 shows a high activity of ZEB2, a promoter of epithelial to mesenchymal transition (EMT), metastasis and resistance in LUAD and breast cancer (Duan et al. 2016; Cui et al. 2019; M.-Z. Li et al. 2017). Coherently, MF2C activity is also inhibited in cluster 5. MF2C is known to be specifically inhibited by miR-223 (secreted by stromal cells) to increase invasion and migration of LUAD (Alečković and Kang 2015).

Cluster 4 (over-represented with colorectal cancer (COREAD)) shows a high activity of BCL6, a novel potential therapeutic target of COREAD (Sun et al. 2020). The role of other top dysregulated kinase and TFs in cluster 4 (MAX, MNT, NCOR2 and UBTF) seem to not have been characterised yet in COREAD, but they have known roles in other cancers. Thus, it could be particularly interesting to investigate those further in the future.

Cluster 6 (over-represented in kidney cancer (ccRCC)) is characterised by a down-regulation of SMARCC2 activity. SMARCC2 is a core subunit of the tumor suppressor complex SWI/SNF A. SWI/SNF inhibition is emerging as a mechanism driving tumor development both in murine model and ccRCC patients (Nargund et al. 2017; Agaimy et al. 2018). The other top down-regulated kinase and TFs of cluster 6 (MYOD1, PBX2, PRMD14 and TFAP4) seem to be less studied in the context of ccRCC. In particular, TFAP4 could be a very interesting protein to study to understand different cancer development mechanisms across different cancers, since it is down-regulated in cluster 6 but up-regulated in cluster 3.

Cluster 7 (mainly characterised by CD8- inflamed tissues over-representation) coherently shows a high activation of NFKB1. Interestingly, RUNX1 is also up-regulated, although it usually suppresses inflammation through NFKB1 inhibition (Bellissimo et al. 2020). This could indicate that perturbation of signaling in the tumor micro-environment prevents the effect of RUNX1 to regulate inflammation. JUN is also up-regulated in cluster 7, coherently with its role in regulation of inflammation in epithelial tissues (Schonthaler, Guinea-Viniegra, and Wagner 2011).

Cluster 8 (over-represented with uterus cancer patients) shows an up-regulation of BCL3, MAZ and MTA2 proteins, as opposed to their down-regulation in cluster 2. BCL3 and MTA2 have been both found to be up-regulated in uterus cancer (Lin et al. 2020) while MAZ dysregulation seems to be under-studied in cancer. HOXB13 is down-regulated in cluster 8. HOXB13 is usually associated with invasiveness of endometrial (uterus tissue) tumors (Zhao, Yamashita, and Ishikawa 2005) and this could indicate that those tumor samples are not yet malignant.

**Systematic exploration of mechanistic hypotheses to connect dysregulated kinases and TF in tumor clusters.**

After highlighting the most consistently deregulated kinase and TFs in each cluster, we sought to investigate the mechanistic links that could connect them together. In particular, we searched which kinases could best explain the downstream deregulated TFs. For that we contextualised a PKN obtained from Omnipath using the top deregulated kinases and TFs of each cluster with CARNIVAL (see methods) (Dugourd et al. 2021; Liu et al. 2019). The 8 causal solution networks (1 per cluster) generated had an average of 75 edges (54 s.d.). In each network, we can explore how given kinases are activating/inhibiting downstream signaling proteins. Indeed kinases and TFs are connected by coherent causal interactions through intermediate signaling proteins (TFs and kisses for which there was not enough target measurement/knowledge to estimate their activities as described in previous sections). Thus, the contextualised causal links also help us hypothesize on the potential activity of intermediate nodes (Supplementary Results Fig 4).

In cluster 1, IRF1, RELA and SPI1 appear to be down-regulated as a consequence of the coordinate up-regulation of PRKCQ and down-regulation of GSK3B, through the down-regulation of SRC and JUN. While SRC is often found up-regulated in many aggressive cancer forms, the up-regualtion of its potential upstream inhibitor PRKCQ and the down-regulation of its downstream target RELA indicate that SRC's role in this tumor cluster might have a more ambivalent role (Irby and Yeatman 2000).

In cluster 2, the up-regulation of BHLHE40 and MAZ can be explained by an upstream up-regulation of the PAK2 kinase, the subsequent up-regulation of TP53,  and the coordinated down-regulation of CDK1. PAK2 is commonly found over-expressed and activated in cancer (Ye and Field 2012). Interestingly, PAK2, TP53 and BHLHE40 are all involved in the regulation of the circadian clock (Karantanos et al. 2013). The PAk2/BHLHE40 crosstalk highlighted here can give an interesting insight into the mechanism through which circadian clock genes can be hijacked in cancer (Sahar and Sassone-Corsi 2009).

In cluster 3, while the poor prognosis marker KDM5B wasn’t included in the causal solution network, we found another interesting signaling cascade in the form of SNAI2 inhibition downstream of the HIPK2 kinase. This can be particularly interesting because SNAI2 is another marker of aggressiveness and  chemotherapy resistance in ovarian cancer, especially to cisplatin treatment (Fan et al. 2020), while HIPK2 over-expression is associated with sensitivity to paclitaxel (Z. Li et al. 2010). Paclitaxel and cisplatin can be used in combinaison, but this data indicates that some patients might benefit more from treatment with paclitaxel alone. Furthermore, ovarian tumor patients, especially with high levels of KDM5B have been shown to be more resistant to chemotherapy mix that don’t include paclitaxel (L. Wang et al. 2015).

In cluster 4, the increased activity of NCOR2, BCL6 and MNT can be explained by the upstream activation of MAPK14. Notably, MAPK14, NCOR2 and BLC6 are all well known to promote survival and resistance to stress for tumor cells, while suppressing their growth (Battaglia, Maguire, and Campbell 2010; Grossi et al. 2014; Cardenas et al. 2017). This cascade is also very coherent with the predicted down-regulation of CHUCK, downstream of MAP3K8. CHUCK can inhibit the downstream part of this cascade (NCOR2 and BCL6) and coherently, its own activity has been associated with tumor growth and proliferation(Chavdoula et al. 2019).

In cluster 5, the down-regualtion of RBPJ and SPI1 combined with the up-regulation ZEB2 can be explained by the activation of HDAC1 downstream of CSNK2A2 kinase. The activation of HDAC1 seems to be a very powerful switch for cancer progression (Cao et al. 2017), which is highlighted here by its ability to inhibit senescence pathways on the one hand (through SPI1 inhibition (Delestré et al. 2017)) while promoting metastasis on the other hand (through ZEB2 activation(Delestré et al. 2017)).

In cluster 6, the down-regulation of MYOD1, SMARCC2, TFAP4 and PBX2 can be explained by the combined effect of MAPK1 down-regulation and PRKCZ up-regulation. The MAPK1 down-regulation could be mediated by the downstream inhibition of CREBBP. This is particularly relevant considering that CREBBP inhibition can either negatively or positively regulate proliferation and invasion of different types of kidney cancer cell lines (X. Wang et al. 2017). The network also shows RELA as a potential activated intermediate of PRKZC. RELA is a potential therapeutic target in ccRCC(Peri et al. 2013), and this network give further insight in its possible role in kidney cancer progression, on the one hand through the inhibition of TF4P4, which itself regulates adhesion to extracellular matrix (proteoglycan production)(Ahrens et al. 2020), and on the other hand through the inhibition of PBX2 to prevent apoptosis(Xu et al. 2016).

In cluster 7, we can see that NFKB1, JUN, ETS1, SPI1 and RUNX1 are all tied in together in a cascade. These 5 up-regulated genes are very coherent with the over-representation of inflamed samples in this cluster. The combined action of PRKG1 and CDK5 seems to be facilitating the activation of NFKB1, and the rest of the cascade consequently. Indeed, PRKG1 and CDK5 seem to be mainly acting through a lift of the inhibition of NFKB1, but it is reasonable to expect that NFKB1 should also be activated by other molecular actors that are not captured in the solution network. In particular, NFKB1 is known to be a very important molecular effector of INFG(Pfeffer 2011). Coherently, INFG itself is one of the most important secreted chemokines of CD8 independent immune response in cancer(Pluhar, Pennell, and Olin 2015). Taken together, these evidences draw the picture of a cluster of tumor samples where the anti-tumoral response of the host is particularly active.

In cluster 8, we can see that the activation of BCL3 can be explained as a direct consequence of CDK5 activity. The deregulation of MAZ and HOXB13 can be explained as a consequence of CDK1 activity through CSNK2A1 and YY1. YY1 has been studied as a potential drug target in HPV infection-induced uterus cancer(He et al. 2011). However, it’s role in inhibiting HOXB13 might complicate the picture, as HOXB13 itself can act as an oncogene(Yamashita et al. 2005). Coherently, YY1 has been recently found to have an ambivalent role in tumor progression, and has been shown to have both onco-supressor and oncogene roles, depending on the tumor context(Sarvagalla, Kolapalli, and Vallabhapurapu 2019).

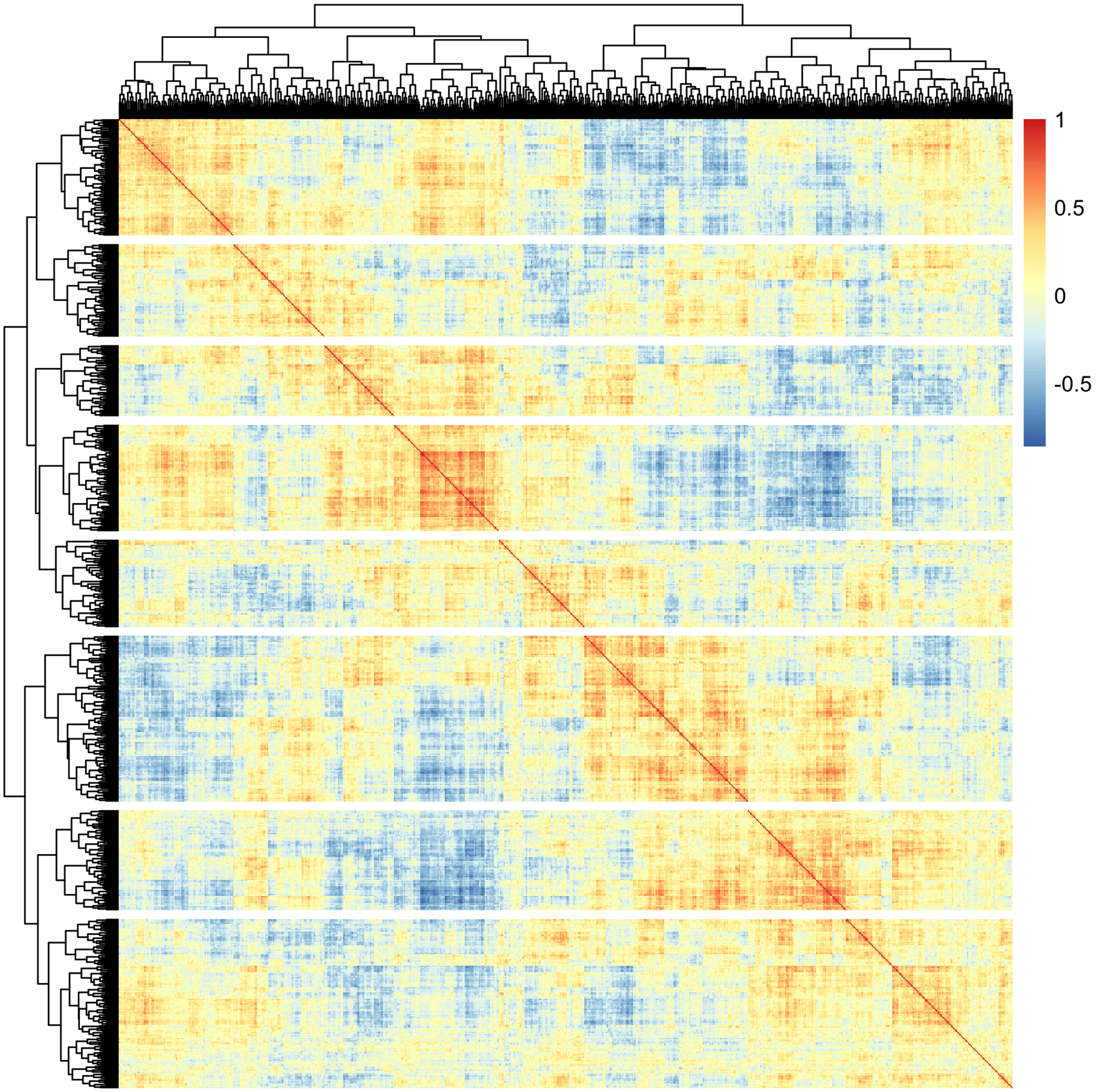


**Supplementary Results Fig 1. Cross-correlation matrix of tumor sample kinase-TF activity signature.** Each cell of the heatmap represents the spearman correlation coefficient between two samples. Correlations are estimated between pairs of vectors combining both TF and kinase activities. Dendrograms represent complete linkage hierarchical clustering from euclidean distances estimated based on the cross-correlation matrix. Each row/column represents a tumor sample.

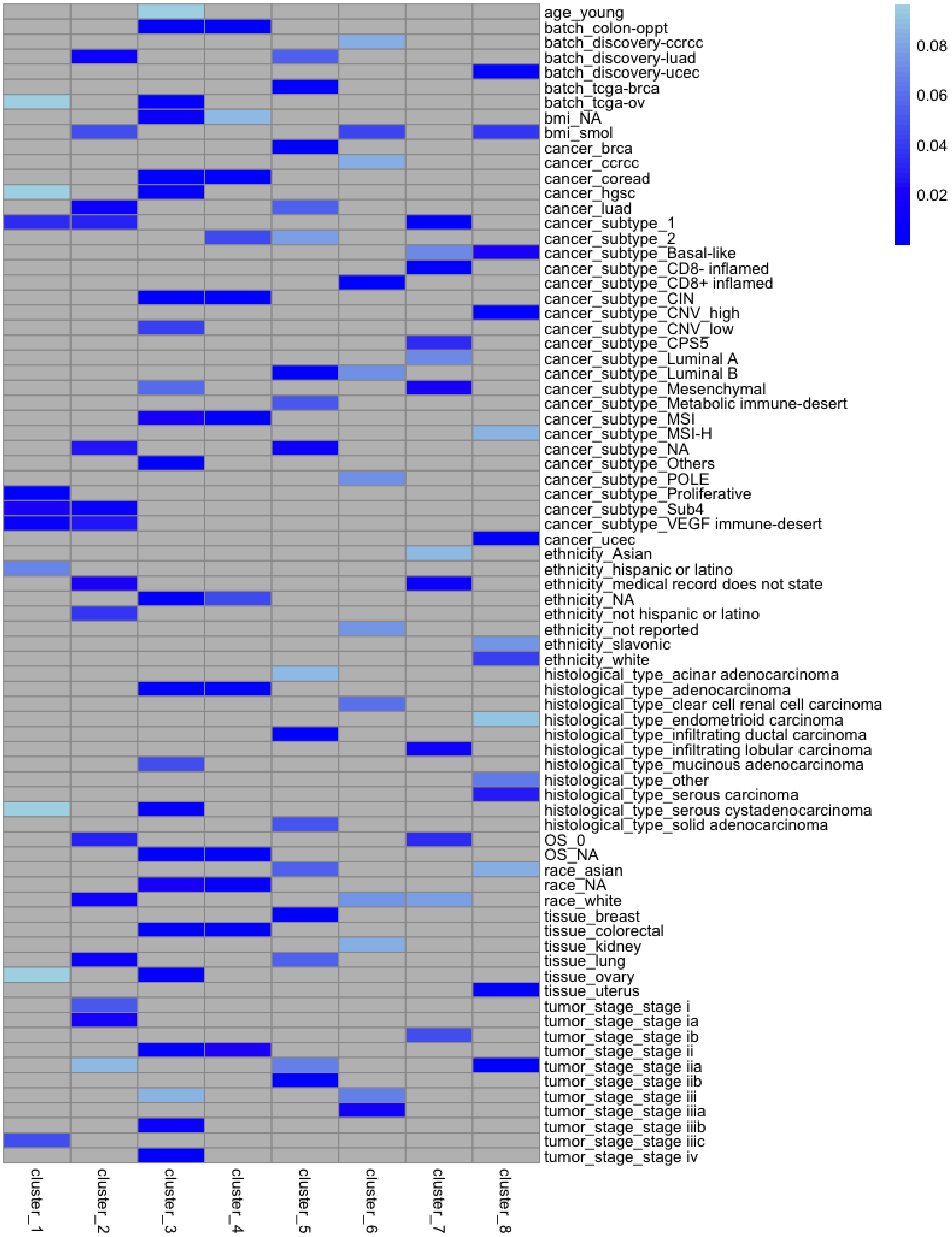


**Supplementary Results Fig 2. heatmap of clinical features over-representation.** Each column represents a given cluster, each row represents a given clinical feature, and each cell represents the p-value of the over-representation of the given clinical feature in the patients of the given cluster. Only p-values < 0.1 are represented with shades of blue, others are greyed out.

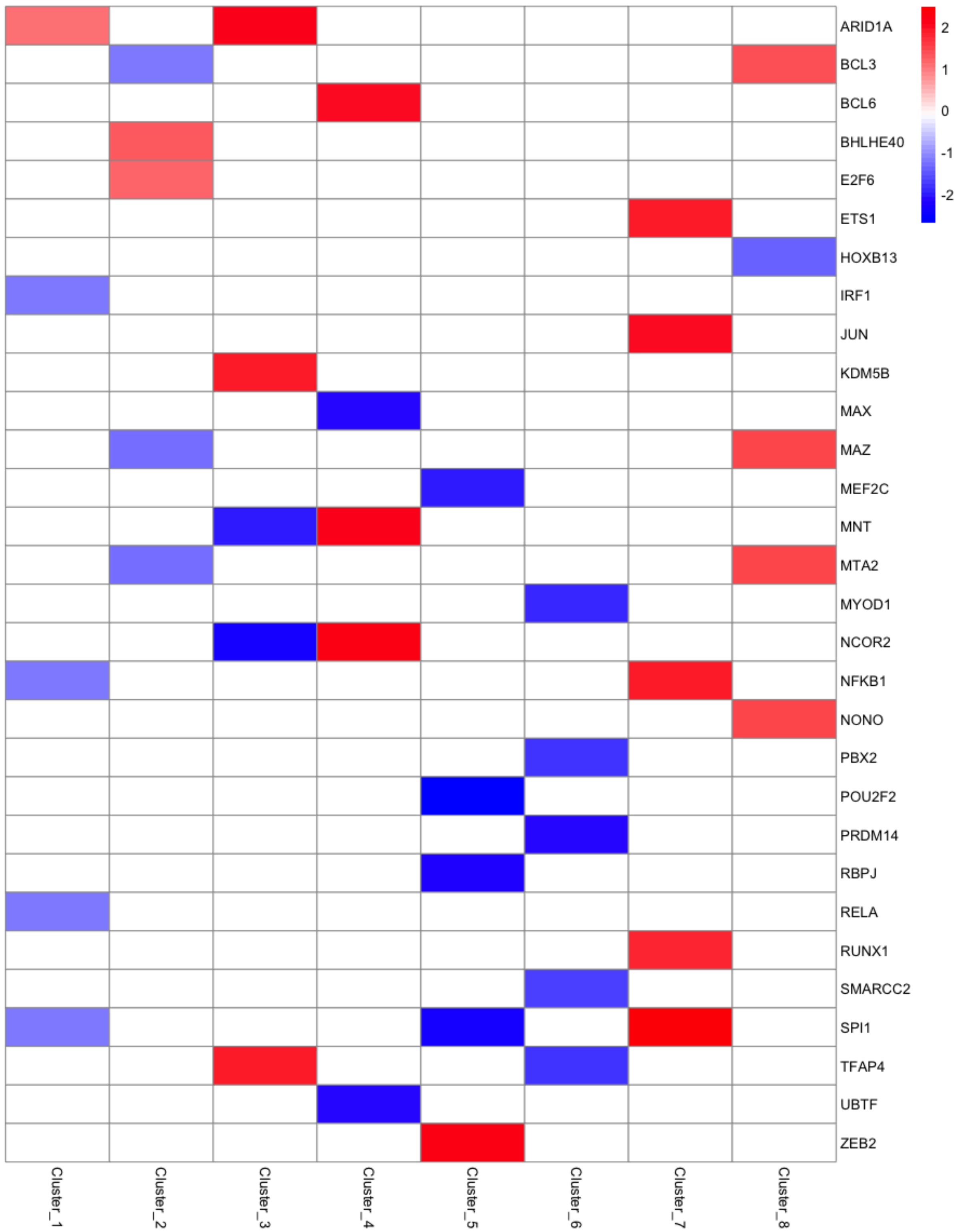

**Supplementary Results Fig 3. Most consistently deregulated kinase and TF activities in each cluster.** Each column represents a given cluster, each row represents a given kinase / TF, and each cell represents the scaled average of the TF/kinase scores across the patients of the given cluster. Only absolute scaled average > 1.7 are displayed as shades of red and blue, others are whited out.

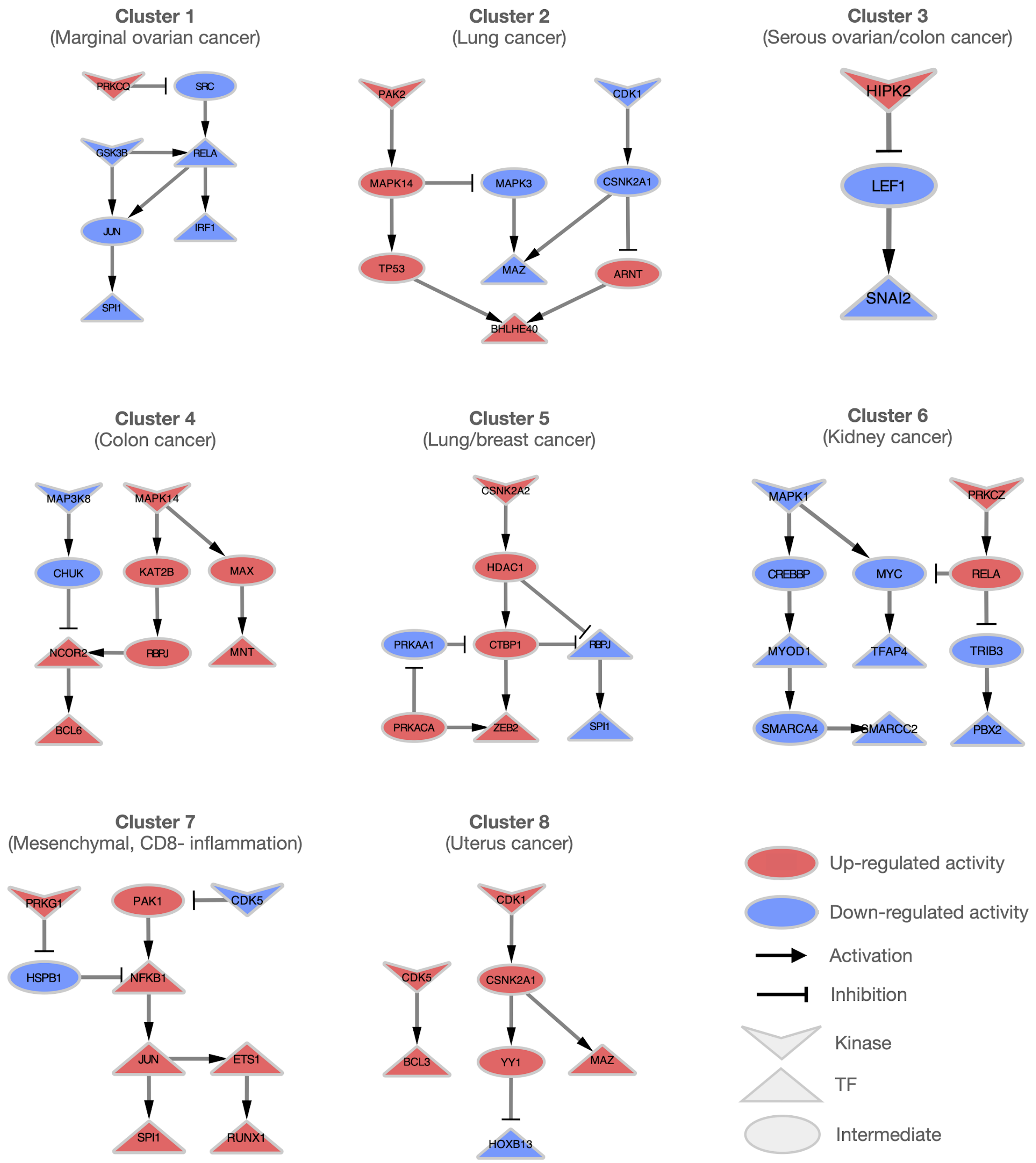

**Supplementary Results Fig 4.** Mechanistic hypotheses to connect the highlighted kinase and TFs of each cancer cluster.

Ahrens, Theresa D., Sara R. Bang-Christensen, Amalie M. Jørgensen, Caroline Løppke, Charlotte B. Spliid, Nicolai T. Sand, Thomas M. Clausen, Ali Salanti, and Mette Ø. Agerbæk. 2020. "The Role of Proteoglycans in Cancer Metastasis and Circulating Tumor Cell Analysis." *Frontiers in Cell and Developmental Biology* 8 (August): 749.

Alečković, Maša, and Yibin Kang. 2015. "Regulation of Cancer Metastasis by Cell-Free miRNAs." *Biochimica et Biophysica Acta* 1855 (1): 24–42.

Battaglia, Sebastiano, Orla Maguire, and Moray J. Campbell. 2010. "Transcription Factor Co-Repressors in Cancer Biology: Roles and Targeting." *International Journal of Cancer. Journal International Du Cancer* 126 (11): 2511–19.

Bellissimo, Dana C., Chia-Hui Chen, Qin Zhu, Sumedha Bagga, Chung-Tsai Lee, Bing He, Gerald B. Wertheim, et al. 2020. "Runx1 Negatively Regulates Inflammatory Cytokine Production by Neutrophils in Response to Toll-like Receptor Signaling." *Blood Advances* 4 (6): 1145–58.

Cao, Lin-Lin, Xiaoxu Song, Lin Pei, Lianhua Liu, Hui Wang, and Mei Jia. 2017. "Histone Deacetylase HDAC1 Expression Correlates with the Progression and Prognosis of Lung Cancer: A Meta-Analysis." *Medicine* 96 (31): e7663.

Cardenas, Mariano G., Erin Oswald, Wenbo Yu, Fengtian Xue, Alexander D. MacKerell Jr, and Ari M. Melnick. 2017. "The Expanding Role of the BCL6 Oncoprotein as a Cancer Therapeutic Target." *Clinical Cancer Research: An Official Journal of the American Association for Cancer Research* 23 (4): 885–93.

Chavdoula, Evangelia, David M. Habiel, Eugenia Roupakia, Georgios S. Markopoulos, Eleni Vasilaki, Antonis Kokkalis, Alexander P. Polyzos, et al. 2019. "CHUK/IKK-α Loss in Lung Epithelial Cells Enhances NSCLC Growth Associated with HIF up-Regulation." *Life Science Alliance* 2 (6). https://doi.org/10.26508/lsa.201900460.

Cohen, Samantha, Rebecca Mosig, Erin Moshier, Elena Pereira, Jamal Rahaman, Monica Prasad-Hayes, Richard Halpert, Jean-Noel Billaud, Peter Dottino, and John A. Martignetti. 2014. "Interferon Regulatory Factor 1 Is an Independent Predictor of Platinum Resistance and Survival in High-Grade Serous Ovarian Carcinoma." *Gynecologic Oncology* 134 (3): 591–98.

Cui, Jianying, Guodong Pan, Qingjuan He, Lizhi Yin, Rui Guo, and Hongmei Bi. 2019. "MicroRNA-545 Targets ZEB2 to Inhibit the Development of Non-Small Cell Lung Cancer by Inactivating Wnt/β-Catenin Pathway." *Oncology Letters* 18 (3): 2931–38.

Delestré, Laure, Hengxiang Cui, Michela Esposito, Cyril Quiveron, Elena Mylonas, Virginie Penard-Lacronique, Oliver Bischof, and Christel Guillouf. 2017. "Senescence Is a Spi1-Induced Anti-Proliferative Mechanism in Primary Hematopoietic Cells." *Haematologica* 102 (11): 1850–60.

Duan, Xunhuang, Zhaojian Fu, Lingyuan Gao, Jin Zhou, Xiaojie Deng, Xiaojun Luo, Weiyi Fang, and Rongcheng Luo. 2016. “Direct Interaction between miR-203 and ZEB2 Suppresses Epithelial-Mesenchymal Transition Signaling and Reduces Lung Adenocarcinoma Chemoresistance.” *Acta Biochimica et Biophysica Sinica* 48 (11): 1042–49.

Dugourd, Aurelien, Christoph Kuppe, Marco Sciacovelli, Enio Gjerga, Attila Gabor, Kristina B. Emdal, Vitor Vieira, et al. 2021. “Causal Integration of Multi-Omics Data with Prior Knowledge to Generate Mechanistic Hypotheses.” *Molecular Systems Biology* 17 (1): e9730.

Fan, Lili, Han Lei, Sai Zhang, Yulong Peng, Chunyan Fu, Guang Shu, and Gang Yin. 2020. “Non-Canonical Signaling Pathway of SNAI2 Induces EMT in Ovarian Cancer Cells by Suppressing miR-222-3p Transcription and Upregulating PDCD10.” *Theranostics* 10 (13): 5895–5913.

Grossi, Valentina, Alessia Peserico, Tugsan Tezil, and Cristiano Simone. 2014. “p38α MAPK Pathway: A Key Factor in Colorectal Cancer Therapy and Chemoresistance.” *World Journal of Gastroenterology: WJG* 20 (29): 9744–58.

He, Guifen, Qian Wang, Yuqi Zhou, Xiaohua Wu, Lan Wang, Nadire Duru, Xiangtao Kong, et al. 2011. “YY1 Is a Novel Potential Therapeutic Target for the Treatment of HPV Infection-Induced Cervical Cancer by Arsenic Trioxide.” *International Journal of Gynecological Cancer: Official Journal of the International Gynecological Cancer Society* 21 (6): 1097–1104.

Irby, R. B., and T. J. Yeatman. 2000. “Role of Src Expression and Activation in Human Cancer.” *Oncogene* 19 (49): 5636–42.

Karantanos, Theodoros, George Theodoropoulos, Maria Gazouli, Anna Vaiopoulou, Christina Karantanou, Maria Lymberi, and Dimitrios Pektasides. 2013. “Expression of Clock Genes in Patients with Colorectal Cancer.” *The International Journal of Biological Markers* 28 (3): 280–85.

Kiss, Zsofia, Maria Mudryj, and Paramita M. Ghosh. 2020. “Non-Circadian Aspects of BHLHE40 Cellular Function in Cancer.” *Genes & Cancer* 11 (1-2): 1–19.

Li, Ming-Zhe, Jing-Jing Wang, Shi-Bin Yang, Wen-Feng Li, Long-Bin Xiao, Yu-Long He, and Xin-Ming Song. 2017. “ZEB2 Promotes Tumor Metastasis and Correlates with Poor Prognosis of Human Colorectal Cancer.” *American Journal of Translational Research* 9 (6): 2838–51.

Lin, Chia-Liang, Tsung-Ho Ying, Shun-Fa Yang, Shih-Wei Wang, Shih-Ping Cheng, Jie-Jen Lee, and Yi-Hsien Hsieh. 2020. “Transcriptional Suppression of miR-7 by MTA2 Induces Sp1-Mediated KLK10 Expression and Metastasis of Cervical Cancer.” *Molecular Therapy. Nucleic Acids* 20 (June): 699–710.

Liu, Anika, Panuwat Trairatphisan, Enio Gjerga, Athanasios Didangelos, Jonathan Barratt, and Julio Saez-Rodriguez. 2019. “From Expression Footprints to Causal Pathways: Contextualizing Large Signaling Networks with CARNIVAL.” *NPJ Systems Biology and Applications* 5 (November): 40.

Li, Zhimin, Sha Hu, Jing Wang, Jing Cai, Lan Xiao, Lili Yu, and Zehua Wang. 2010. “MiR-27a Modulates MDR1/P-Glycoprotein Expression by Targeting HIPK2 in Human Ovarian Cancer Cells.” *Gynecologic Oncology* 119 (1): 125–30.

Malvi, Parmanand, Radoslav Janostiak, Arvindhan Nagarajan, Guoping Cai, and Narendra Wajapeyee. 2019. “Loss of Thymidine Kinase 1 Inhibits Lung Cancer

Growth and Metastatic Attributes by Reducing GDF15 Expression." *PLoS Genetics* 15 (10): e1008439.

Nargund, Amrita M., Hatice U. Osmanbeyoglu, Emily H. Cheng, and James J. Hsieh. 2017. "SWI/SNF Tumor Suppressor Gene PBRM1/BAF180 in Human Clear Cell Kidney Cancer." *Molecular & Cellular Oncology* 4 (4): e1342747.

O'Donnell, Amanda J. M., Kenneth G. Macleod, David J. Burns, John F. Smyth, and Simon P. Langdon. 2005. "Estrogen Receptor-Alpha Mediates Gene Expression Changes and Growth Response in Ovarian Cancer Cells Exposed to Estrogen." *Endocrine-Related Cancer* 12 (4): 851–66.

Peri, Suraj, Karthik Devarajan, Dong-Hua Yang, Alfred G. Knudson, and Siddharth Balachandran. 2013. "Meta-Analysis Identifies NF-κB as a Therapeutic Target in Renal Cancer." *PloS One* 8 (10): e76746.

Pfeffer, Lawrence M. 2011. "The Role of Nuclear Factor κB in the Interferon Response." *Journal of Interferon & Cytokine Research: The Official Journal of the International Society for Interferon and Cytokine Research* 31 (7): 553–59.

Pluhar, G. Elizabeth, Christopher A. Pennell, and Michael R. Olin. 2015. "$CD8^{+}$ T Cell-Independent Immune-Mediated Mechanisms of Anti-Tumor Activity." *Critical Reviews in Immunology* 35 (2): 153–72.

Sahar, Saurabh, and Paolo Sassone-Corsi. 2009. "Metabolism and Cancer: The Circadian Clock Connection." *Nature Reviews. Cancer* 9 (12): 886–96.

Sarvagalla, Sailu, Srinivasa Prasad Kolapalli, and Sivakumar Vallabhapurapu. 2019. "The Two Sides of YY1 in Cancer: A Friend and a Foe." *Frontiers in Oncology* 9 (November): 1230.

Schonthaler, Helia B., Juan Guinea-Viniegra, and Erwin F. Wagner. 2011. "Targeting Inflammation by Modulating the Jun/AP-1 Pathway." *Annals of the Rheumatic Diseases* 70 Suppl 1 (March): i109–12.

Sun, Naihui, Liang Zhang, Chongguang Zhang, and Yuan Yuan. 2020. "miR-144-3p Inhibits Cell Proliferation of Colorectal Cancer Cells by Targeting BCL6 via Inhibition of Wnt/β-Catenin Signaling." *Cellular & Molecular Biology Letters* 25 (March): 19.

Wang, Lishuang, Yuanfu Mao, Guiqin Du, Chunbo He, and Shiyu Han. 2015. "Overexpression of JARID1B Is Associated with Poor Prognosis and Chemotherapy Resistance in Epithelial Ovarian Cancer." *Tumour Biology: The Journal of the International Society for Oncodevelopmental Biology and Medicine* 36 (4): 2465–72.

Wang, Xue, Hui Cui, Zhongguan Lou, Shuaishuai Huang, Yu Ren, Ping Wang, and Guobin Weng. 2017. "Cyclic AMP Responsive Element-Binding Protein Induces Metastatic Renal Cell Carcinoma by Mediating the Expression of Matrix Metallopeptidase-2/9 and Proteins Associated with Epithelial-Mesenchymal Transition." *Molecular Medicine Reports* 15 (6): 4191–98.

Xu, Chengshan, Hengheng Li, Ling Zhang, Tianjun Jia, Lianning Duan, and Chengrong Lu. 2016. "MicroRNA‑1915‑3p Prevents the Apoptosis of Lung Cancer Cells by Downregulating DRG2 and PBX2." *Molecular Medicine Reports* 13 (1): 505–12.

Yachida, Nozomi, Kosuke Yoshihara, Kazuaki Suda, Hirofumi Nakaoka, Haruka Ueda, Kentaro Sugino, Manako Yamaguchi, et al. 2020. "ARID1A Protein Expression Is Retained in Ovarian Endometriosis with ARID1A Loss-of-Function Mutations:

Implication for the Two-Hit Hypothesis." *Scientific Reports* 10 (1): 14260.
Yamashita, Tsuyoshi, Zhao Yawei, Seisiro Tazawa, Kiichiro Kawai, Hideto Katayama, and Mutsuo Ishikawa. 2005. "Regulation of Tumor Invasion by HOXB13 Gene Overexpressed in Human Endometrial Cancer." *Cancer Research* 65 (9 Supplement): 1328–29.
Ye, Diana Zi, and Jeffrey Field. 2012. "PAK Signaling in Cancer." *Cellular Logistics* 2 (2): 105–16.
Zhang, Bin, Hao Zhang, and Gang Shen. 2015. "Metastasis-Associated Protein 2 (MTA2) Promotes the Metastasis of Non-Small-Cell Lung Cancer through the Inhibition of the Cell Adhesion Molecule Ep-CAM and E-Cadherin." *Japanese Journal of Clinical Oncology* 45 (8): 755–66.
Zhao, Yawei, Tsuyoshi Yamashita, and Mutsuo Ishikawa. 2005. "Regulation of Tumor Invasion by HOXB13 Gene Overexpressed in Human Endometrial Cancer." *Oncology Reports* 13 (4): 721–26.
